# Large-scale assessment of habitat quality and quantity change on declining European butterflies

**DOI:** 10.1101/2022.09.29.510048

**Authors:** Nicolas Chazot, Søren Faurby, Chris van Swaay, Johan Ekroos, Niklas Wahlberg, Christine D. Bacon, Alexandre Antonelli

## Abstract

The rapid decline of biodiversity as a consequence of increased environmental impact by human activities requires urgent action against the ongoing crisis. At the heart of conservation policy is the debate on quality versus quantity: should the post-2020 Global Biodiversity Framework focus on maximising total protected area, or pursue instead qualitative targets? To improve conservation practices, we need to both understand the current trend of biodiversity and the factors driving the decline. We address this by: (1) projecting current European butterfly population trends for the next 50 and 100 years, (2) comparing the decline in species richness, phylogenetic diversity and habitat specialization among different habitats, and (3) estimating the relationship between recent trends in habitat quality or quantity and the decline of multiple metrics of European butterfly diversity. We do not find any significant relationship between diversity loss and habitat area loss and conclude that neither long-term nor short-term changes in habitat area are primary predictors of large-scale butterfly decline in Europe. Habitat quality emerges as the most crucial factor in our analyses – both the area affected and the severity of habitat quality reduction. Habitat degradation causes vegetation changes in structure and diversity, which affect butterfly survival. We also estimate a higher decline of habitat specialists than generalist species. We conclude that habitat protection cannot focus solely on the maximization of habitat area but urgently needs to restore high quality ecosystems to provide the full range of ecological requirements for biodiversity.

## Introduction

According to the recent global assessment from the Intergovernmental Science Policy Platform on Biodiversity and Ecosystem Services (IPBES, 2019), about a million species are likely at risk of extinction. The increasing pressure exerted by human-altered landscapes (such as land cleared for agriculture, urbanization and exploration of natural resources) on biodiversity-rich habitats stresses the urgency of prioritising conservation targets and developing effective conservation practices. In particular, the discussions on area-based conservation targets, such as protecting “30% of the Earth’s land and oceans by 2030” (Woodley et al. 2019, Maxwell et al. 2019, Obura et al. 2021) has played a prominent role in the run-up to the post-2020 Global Biodiversity Framework under the Convention on Biological Diversity, to be set at the next Conference of the Parties on Biodiversity (COP15). Evidence-based assessments of such goals and their relation to more qualitative targets are therefore urgently needed.

The state of terrestrial vertebrate biodiversity and trends over time are relatively well documented and point at an extinction pattern approaching past mass-extinction events (e.g. Ceballos et al. 2015, Andermann et al. 2020, Neubauer et al. 2021). Reports on the state of insect biodiversity have only recently increased in number. Available evidence shows that insects are undergoing declines in terms of both biomass and abundance. Hallmann et al. (2017) reported a seasonal decline of insect biomass of 76% over 27 years across Germany. Seibold et al. (2019) recorded an average of 67%, 78% and 34% decline in the biomass, abundance and number of grassland-inhabiting insect species between 2008 and 2017. In a study focusing on British moths, Macgregor et al. (2019) identified a steady decline between 1982 and 2017. While studies have repeatedly shown that global insect diversity is on the decline, generalizations regarding the magnitude and rate of this decline is challenged by the paucity of information available across most insect groups (Wagner 2020).

Butterflies are amongst the most well-studied insect groups, and reports suggest that many species have greatly declined, at least during the last 50–100 years in Europe (Kuussaari et al. 2007, van Strien et al. 2019) and elsewhere (Schlicht and Orwig 1998). In particular, evidence points to a high sensitivity of poorly mobile species with high habitat specificity (Eskildsen et al. 2015), and recent systematic butterfly monitoring schemes show that e.g. grassland butterflies are still declining (van Swaay et al. 2019, WWF 2020). Following Wiemers et al. (2018), there are 496 species of butterflies in Europe and according to the 2009 report from the International Union for the Conservation of Nature (IUCN), about a third are declining. Both life history traits and external factors, in particular land use changes and climate change explain the decline of butterflies (Wagner, 2020). Thus, identifying the consequences of human activities on butterfly diversity is paramount to change detrimental practices and design efficient conservation strategies, in particular the management of natural and semi-natural ecosystems.

A key challenge in conservation is to identify whether promoting habitat quantity or habitat quality best ensures long-term preservation of biodiversity (e.g. Thomas et al. 2001). On the one hand, maximizing habitat area is critical as it directly benefits local population size of wild species, enables dispersal between patches, and thus contributes to survival of populations at the landscape level (Hanski & Gilpin 1991, Hanksi & Thomas 1994). In local surveys, butterflies with low dispersal ability often decline faster than species with high dispersal ability (e.g., Maes and van Dyck 2001, Filz et al. 2013). Therefore, in addition to habitat area, the distance and complementarity between patches will determine the survival of organisms occurring in fragmented landscapes. On the other hand, small changes in habitat state or quality within a single patch can lead to rapid changes in the community. Shifts in plant community composition as a result of landscape changes can impact on vegetation structure, microclimate and host-plant availability (e.g. Stefanescu et al. 2009, Maes and van Dyck, 2001). For individual butterfly species, habitat quality is defined in terms of resource availability, such as host-plant abundance and occurrence of nectar-producing flowers. Because species differ in resource use, defining habitat quality for entire communities is not trivial. Nevertheless, maintaining diverse plant communities through the preservation of high-quality habitat patches provides a diversity of resources fundamental for the conservation of local butterfly diversity.

While it is clear that both habitat quantity and quality play a role in the maintenance of butterfly diversity, most attempts to compare their relative importance have focused on the local scale, following population changes over time within sites or at the metapopulation scale for small subsets of species (e.g. Dennis & Eales 1997, Thomas et al. 2001). A local scale approach allows for a fine-grain estimation of population trends and a direct assessment of the relationship between changes in habitats, community composition, vegetation and species traits. However, generalizing these local scale results to identifying overarching patterns requires many of such studies to produce comprehensive sets of information. Instead, here we focus on large scale assessments of populations and habitats to investigate the relative effects of changes in habitat quantity and quality on butterfly trends on the European continent (Wiemers et al. 2018). To address this question, we first projected the relative population sizes in the future from butterfly population trends estimated at the European scale between 1999 and 2009. Using a dataset of relative frequency of habitat use, we assessed changes in community composition with three different metrics: species richness, phylogenetic diversity and community specialization. We then compiled average estimates of habitat changes across Europe, both in terms of quantity and quality. Finally, we tested which habitat variables best predict the future changes in species composition across habitats.

## Materials and methods

### Data

Butterfly population trends – To predict future population change, we used the population trends estimated for establishing the European Red List of butterflies in 2009 (Swaay et al 2010); to our knowledge the most recent complete assessment of trends for Europe. These trends were derived by combining national trend estimates provided by national experts, into European scale estimations. They inform about population dynamics as percentages of changes between 1999 and 2009 for all species of European butterflies as estimated by specialists at the scale of Europe. Each trend had a mean and a lower and higher estimation bound (Supplementary Table S1.1).

Butterfly habitat use – To compare the changes in biodiversity among habitats, we assigned species of butterflies to 29 habitat categories based on earlier assessments (van Swaay et al. 2006). In this dataset, habitat use was scored by 50 specialists across Europe (Supplementary Table S1.2). Habitat classification used the Corine biotope system (based on Moss et al 1991). Instead of coding the presence or absence of species in each habitat, we transformed the number of times a species was scored in a habitat into percentages of relative habitat occupancy. In this way, we obtained an estimation of the relative importance of a habitat in the ecology of a butterfly species and therefore a relative frequency of habitat use for each species.

Phylogenetic tree of European butterflies – To estimate future changes in phylogenetic diversity, we used the complete phylogenetic tree of European butterflies from Wiemers et al. (2020). We extracted 1000 random time-calibrated trees from the posterior distribution to account for phylogenetic uncertainty.

Trends in habitat quantity – We used two published sources of information to obtain trends in habitat cover: the European Red List of habitats (Janssen et al. 2016) and the Corine Land Cover dataset (https://land.copernicus.eu/pan-european/corine-land-cover, version V2018_20).

First, we used the net percentage of habitat reduction over the last 50 years reported for the different habitats in the European Red List of habitats. We recovered the amount of surface loss from the estimated current area of distribution reported in the same documents. For eight habitats out of 170, either the information about current area of distribution or a quantitative trend was not available; those habitats were removed from the analysis.

Second, we used data from the Corine Land Cover European program to estimate trends between 2000 and 2006 (https://www.eea.europa.eu/data-and-maps/data/corine-biotopes). We downloaded the Land Cover Change 2000-2006 dataset, whose time interval overlaps with the butterfly population trends estimated between 1999 and 2009. For each habitat, we defined habitat gain and habitat loss by calculating the number of cells (at 25-ha resolution) lost or gained during the time period considered for land-cover classes in the CORINE databases (Bossard et al. 2000). We then compared these to the Corine Land Cover data in 2000 to obtain percentages of changes. To determine the net change in habitat cover across Europe, we calculated the difference between the percentage of cells gained and lost for each biotope. In order to match the data available for the butterfly population trends we removed data for Russia, Ukraine, Belarus and the Vatican. The Red List dataset, the Land Cover Change dataset and the one used for classifying butterflies did not match: the European Red List of habitats uses the EUNIS classification system, whereas the Land Cover Change uses a Corine classification slightly different from the Corine classification used for the butterfly dataset. Hence, we used two correspondence tables to link the Red List to Land Cover Change classification (EUNIS/ Land Cover Change equivalence table provided by the European Environment Agency) (https://www.eea.europa.eu/data-and-maps/data/eunis-habitat-classification/documentation/eunis-clc.pdf) and from the Land Cover Change to the butterfly classification system (our own equivalence table). In cases where one biotope in a dataset corresponded to several biotopes from the other dataset, we split the amount of changes equally among the corresponding biotopes (Supplementary Table S1.3, S1.4).

Trends in habitat quality – To obtain trends in habitat quality at the European scale we used the estimations of biotic and abiotic changes as reported in the European Red List of habitats (Janssen et al. 2016). For each biotope we recovered the extent (percentage) of the area affected by reduction in abiotic and/or biotic quality and the estimated relative severity of the reduction as estimated by Janssen et al. (2016). We transformed the percentages into surface area using the estimated current area of occupancy, available from the same data source. For 27 habitats, quality trend information was not available. These represented about 1.7% of the total surface covered by our dataset and were removed from the analysis. We also combined the area and severity variables by multiplying one with the other (hereafter areaXseverity) to test whether the combination of both variables was a better predictor than each variable separately.

### Analyses

We aimed at estimating the changes in European butterfly communities in 50 and 100 years (considering the baseline year of 2019) resulting from the declining species if they follow population trends estimated between 1999-2009. We projected future populations based on these estimations for the period between 1999-2009. We combined this information with data on relative habitat use to estimate potential diversity loss in different habitats. This projected diversity loss was finally compared with estimated changes in habitat quantity and quality in the past to assess which of the two best predicts the decline of diversity in the different habitats (Fig. 1).

**Figure 1.**
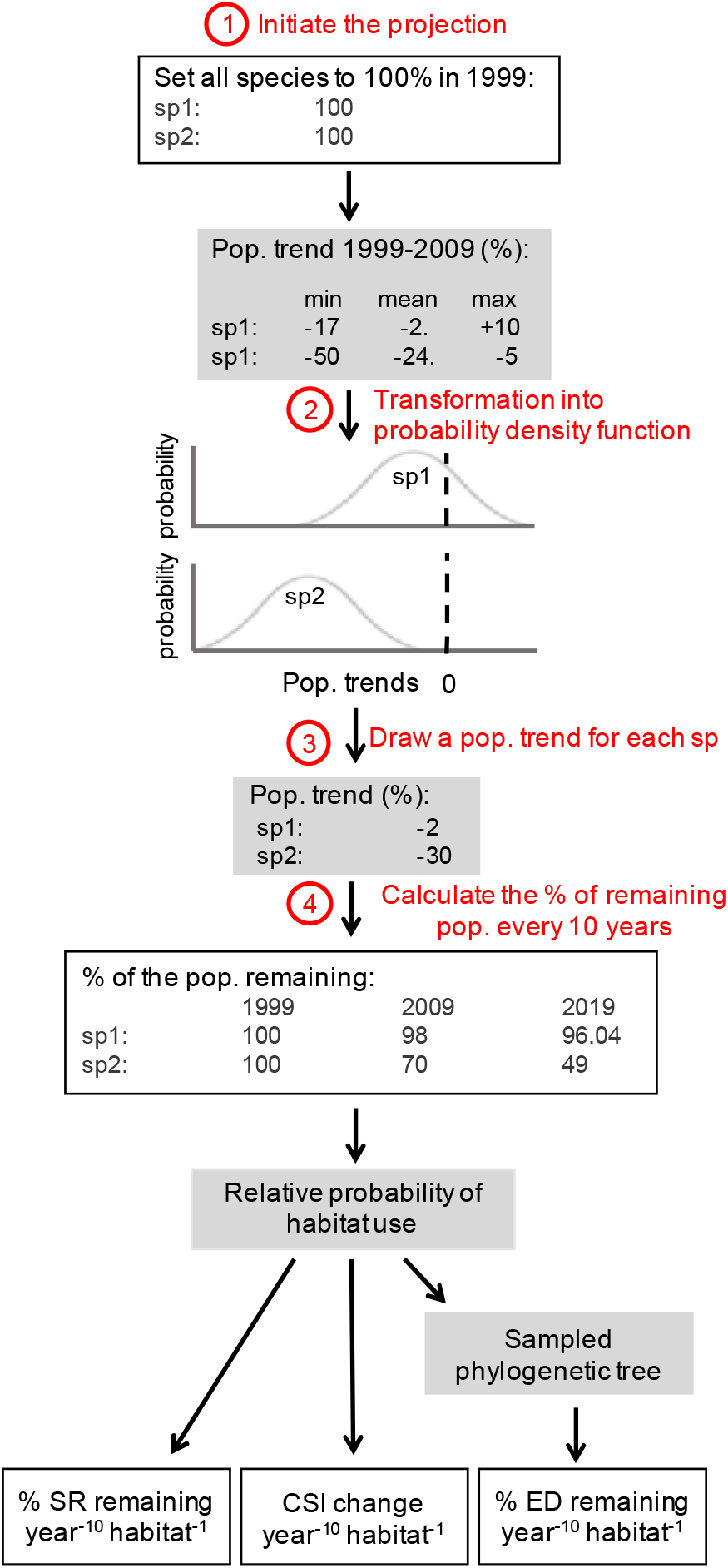
Schematic representation of the different steps performed for a single projection. This procedure was repeated 100 times to obtain 100 projections. We set all species populations at 100% in 1999 to initiate the projection (1). For each species, we fitted a distribution to the population trends information available (2). We sampled a population trend for each species within its distribution (3). Then, at every time t in the future (every 10 years, until 2119), we applied the sampled population trend to the population state at t-1 to calculate the percentage of the initial population remaining (4). We sampled one phylogenetic tree from the posterior distribution of European butterfly tree and used that tree, as well as habitat use information to calculate diversity indices: SR, ED and CSI. We repeated that procedure 1000 times.

Future projections – We calculated the future state of diversity based on the relative population declines reported between 1999 and 2009, assuming (for the sake of the models) populations to be at 100% in 1999 (Fig. 1). We sampled a percentage of decline of each species in a normal function fitted to the minimum, maximum and mean trend of each species. Because we assumed the population trends to be normally distributed between the minimum and maximum values available, we approximated the standard deviation of the normal function with (maximum -minimum)/4. Every 10 years (population trends available were estimated between 1999 and 2009), we computed the percentage of the initial population remaining until 2119. We repeated these projections 1000 times, sampling a new trend for each species at every new iteration. We report and discuss the results of decline in 2019 plus 50 years and 2019 plus 100 years. At every point in time *t*, the state of a population was calculated by removing a percentage of the population at its state *t-1*. Importantly, we decided to focus on declining species only at each iteration. Increasing population trends were considered as stable. This choice was justified to focus on assessing the importance of habitat changes on negative trends specifically and to avoid community changes to be masked by few exponentially increasing populations.

Species richness (SR) – For each habitat, every 10 years, we calculated the remaining species richness weighted by both the percentage of remaining total population and the relative frequency of habitat use by each species.

Evolutionary Distinctiveness (ED) – We calculated the decline of phylogenetic diversity using Evolutionary Distinctiveness metrics (Isaac et al. 2007). To be able to compare the ED loss through time and across the different habitats, the ED score of each species at each point in time was calculated by weighting the length of the phylogenetic branches by the percentage of remaining population and the relative frequency of habitat use (Supplementary Information S2). Thus, for a single species, the sum of weighted ED scores across all habitats occupied by that species was equal to the standard unweighted ED score. As such, a species able to occupy multiple habitats (‘generalists’ in our terminology) did not contribute multiple times equally to the total ED of different habitats, but instead contributed proportionally to its use of the habitat and its remaining population. For each habitat and every 10 years, we calculated the total ED as the sum of the weighted ED of all species occupying that habitat. We calculated the decline of ED for 100 phylogenetic trees from the posterior distribution of trees, sampling for each tree a different projection.

Null projections – We generated scenarios in which species-specific population trends were permuted among species before projections, to assess whether some habitats are declining slower or faster than a scenario where population trends are not correlated with habitats. We performed 100 additional projections after permutation, and calculated species richness and evolutionary distinctiveness loss under these scenarios. Thus, we compared predicted biodiversity loss in each habitat to the expectation under permuted population trends.

Community Specialisation Index (CSI) – We tested whether the changes in species richness were also accompanied by changes in the relative proportion of habitat specialists and generalists. We used the Community Specialisation Index (CSI) to characterize the community (Devictor et al. 2008). The CSI of a community is the sum of each species specialization index (SSI, Fridley et al. 2007), weighted by the species abundance in the community. SSI is the specialisation index of each species and is calculated as the species density coefficient of variation (standard deviation divided by the mean). We calculated the SSI from the dataset of relative frequency of habitat use. Then we calculated the CSI of each habitat, weighting the SSI by the percentage of remaining population of each species in 1999 (starting year for population trends), also in 50 and 100 years from present day (i.e. 2069 and 2119) for 1000 projections. CSI increases with an increasing proportion of specialists in the community. If species richness loss in a habitat is homogeneously distributed across species, CSI remains identical through time. Any departure from the CSI score in 1999 indicates a change in the relative proportion of specialists and generalists. An increase of CSI through time would indicate an increasing proportion of habitat specialists, hence generalist species declining faster. A decreasing CSI through time would indicate a decreasing proportion of habitat generalists, hence specialist species declining faster.

Linear regressions – To test for relationships between habitat changes and diversity changes we fitted linear models between the habitat trends, total habitat area, and the different metrics for diversity changes in each habitat between 1999 and 2069 (50 years from present day) and between 1999 and 2119 (100 years from present day), including SR, ED, and CSI. To carry out theses analyses we reduced the dataset to a set of habitats for which all trend variables (population and habitats) were available and contained at least three butterfly species. This resulted in a dataset of 23 habitats: three heath and scrub habitats, five grasslands habitats, four forest habitats, four wetland habitats, two unvegetated habitats, three agricultural habitats and one urban habitat (Supplementary Information S1-S3). Linear models were ranked according to their AIC scores. We also tested whether habitat CSI in 1999 predicted the amount of SR and ED lost in 2069 and 2119 using linear regressions.

## Results

### Population trends and predicted diversity loss

The biotopes projected to suffer the highest loss of butterfly species include Bogs And Mires, Unimproved Semi-Natural Grasslands, and Coniferous Woodlands. In contrast, the biotopes with the lowest projected loss of butterflies include Coastal Sand Dunes and Sandy Beaches, Cliffs And Rocky Shores, Shrubland, Arable Land, Urban parks And Fallow Land (Supplementary Information S3). Concerning Wetland biotopes, in which butterfly diversity is the most endangered according to our projections, we estimate an average loss of 39 % [95% CI = 30-47] of butterfly species richness by 2069 and 50 % [40-60] by 2119, based on the rate of species-specific population declines estimated between 1999 and 2009. Overall, we estimate the species richness of butterflies to decline by 24 % [18-31] in Europe by 2069, and 32 % [24-41] by 2119 according to the population trends estimated between 1999 and 2009. Importantly, as stated before, these trends are estimated without accounting for increasing populations thus only estimating changes resulting from declining species. We remain cautious with these estimations of absolute changes and primarily focus on the relative losses between habitats.

Overall, trends projected for evolutionary distinctiveness (ED) are very similar to those of SR when classified by biotope (Supplementary Information S3). Only in the case of Towns, Villages And Industrial Sites it is predicted that phylogenetic loss will be substantially lower than species loss. When averaged across all habitats, the estimated mean ED loss is 25 % [18-33] and 32.8 % [25-44] in 2069 and 2199 respectively.

The Community Specialisation Index (CSI) in 1999 is lowest for the three Agricultural habitats and highest for Sclerophylous Shrub, Broadleaved Evergreen Woodlands, Alpine And Subalpine Grasslands. Of the 22 habitats, we find 16 habitats with decreasing CSI by 2069, indicating a disproportional loss of habitat specialist butterflies across the majority of habitats (Supplementary Information S4). CSI increased for four habitats by 2069 (Sclerophylous Shrub, Coniferous Woodlands, Crops, Heath And Scrubs). CSI almost does not change for two habitats (Improved Grasslands, Dry Calcareous Grasslands And Steppes). Broadleaved Evergreen Woodlands, Alpine And Subalpine Grasslands (two of the most specialized habitats) are characterized by the strongest decline in habitat specialization. Overall, the results remained the same for CSI change between 1999 and 2119 (Supplementary Information S4).

### Predicted diversity loss and habitat changes

Across all 23 biotopes, we find no significant relationship between predicted SR loss or ED loss and any measure of habitat quality or quantity changes (Supplementary Information S5-S7, Fig. 2, 3). However, Fig. 2 and Fig.3 show that habitats belonging to the same higher-level classification are generally very homogeneous, especially for habitat quality changes. Thus, we tested the hypothesis that one group of habitat behaves as an outlier of relationships. We focused on the three most frequent higher-level classes of habitat: wetlands, forests, grasslands. We removed either all forest habitats, all grasslands habitats or all wetland habitats, and fitted again linear models. We find that predicted diversity loss (SR and ED) is significantly higher with greater area affected by habitat quality change when removing all four forest habitats (Supplementary Information S7, Fig. 2, 3). Forest habitats clearly behave as outliers with a much lower diversity loss than predicted by the relationship with habitat quality change area. By removing all forest habitats, we also find the area × severity-index to significantly predict diversity loss, but only for ED loss (Fig. 3). Second, we find a positive relationship between the severity of habitat quality degradation and predicted SR loss or ED loss when all four wetland habitats are removed (Supplementary Information S7). Wetland habitats show lower diversity loss than predicted by the relationship with severity of habitat quality degradation (Fig. 2, 3). Removing either Forest habitats or Wetland habitats does not change any of the relationships with habitat quantity changes. All these results are identical for the predicted diversity loss in 2069 and 2119 (Supplementary Information S7).

**Figure 2.**
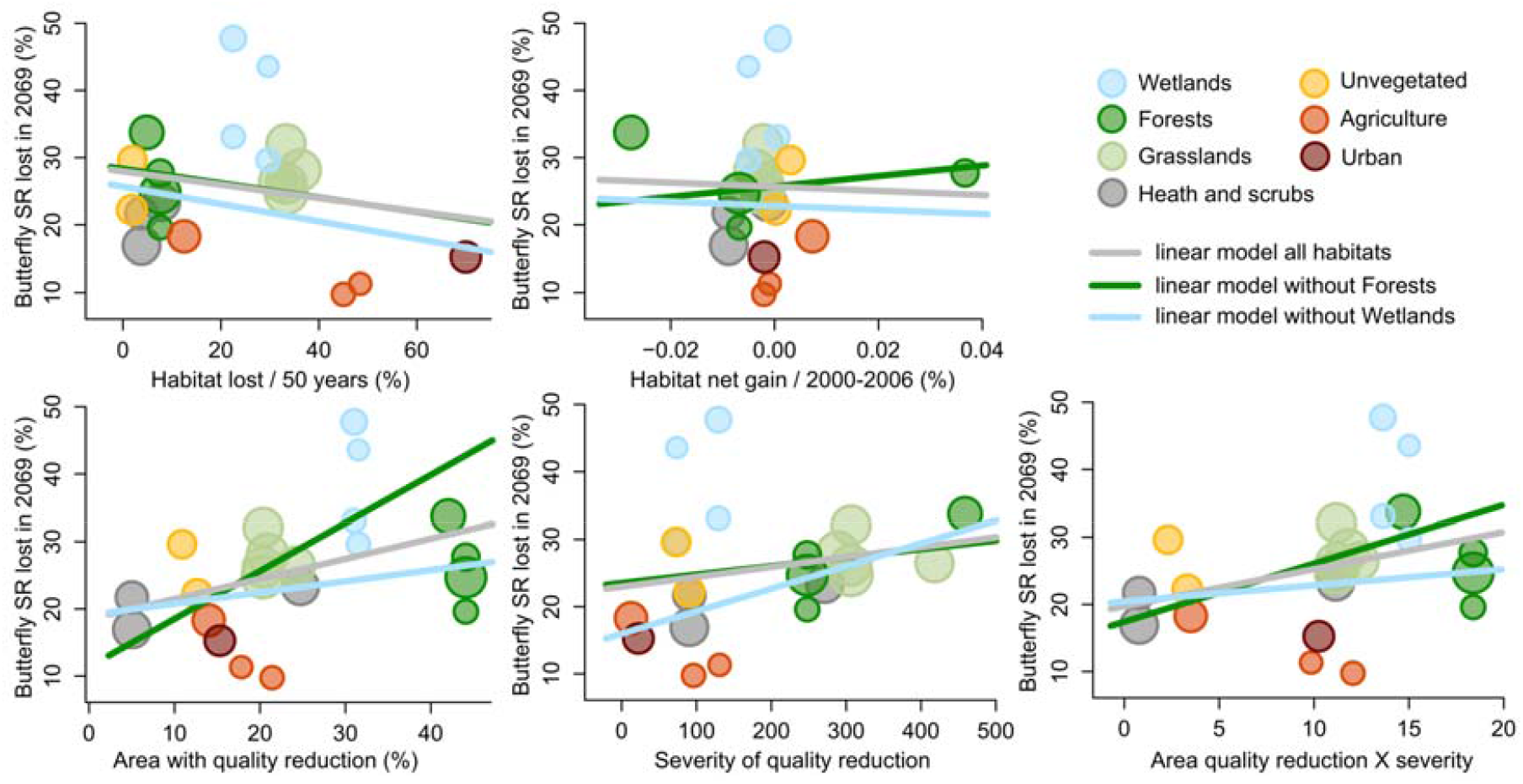
Relationship between the predicted species richness (SR) lost in 2069 and habitat net gain over the last 50 years, habitat net gain between 2000 and 2006, extent of habitat quality reduction, severity of habitat quality reduction, and the product of area loss and the severity of habitat quality loss. Each point is a biotope, coloured according to the higher biotope classification. Point size is proportional to the number of species in each biotope.

**Figure 3.**
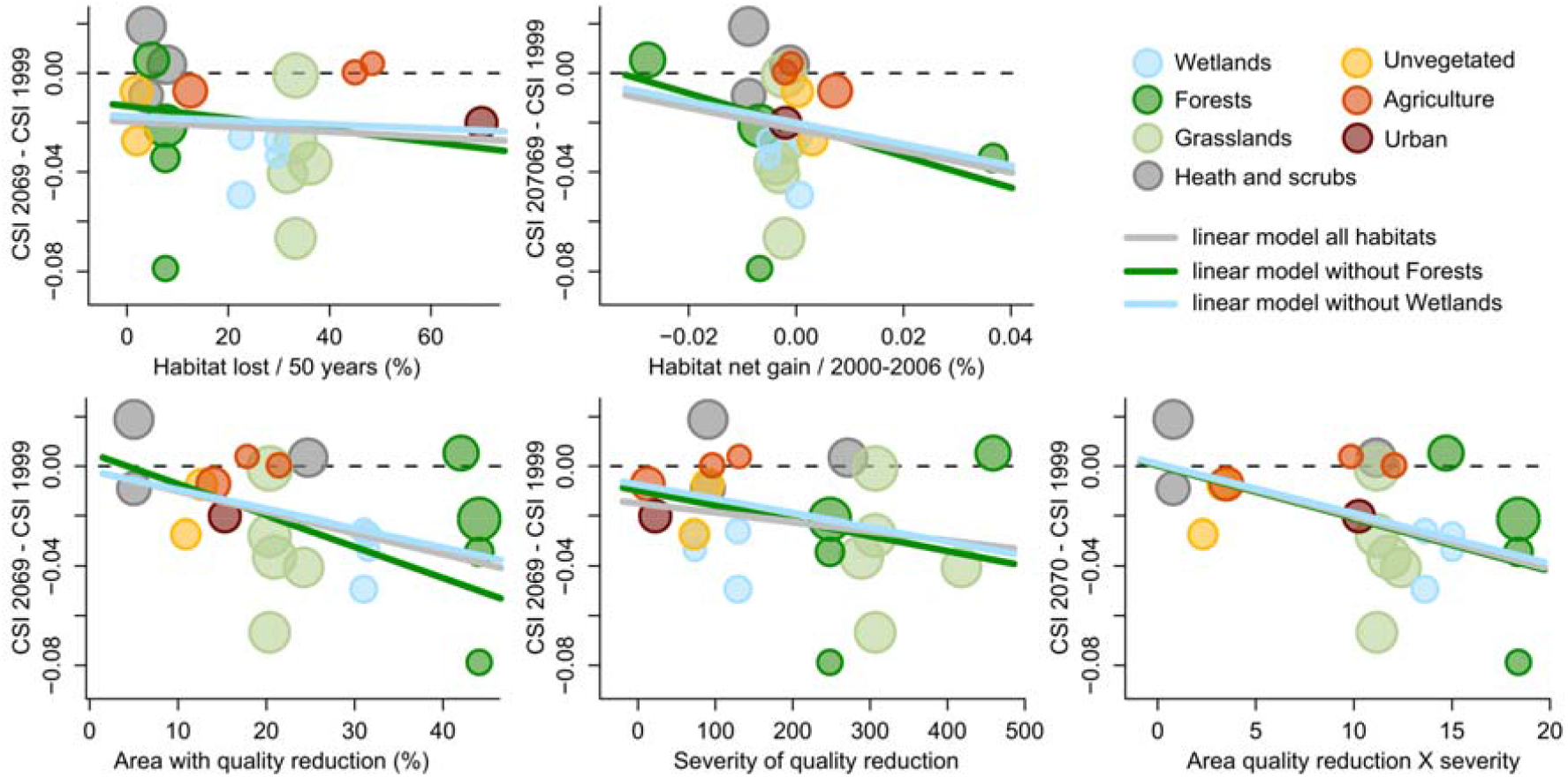
Relationship between the predicted community specialization index (CSI) change in 2069 (CSI 2069 minus CSI 1999) and habitat net gain over the last 50 years, habitat net gain between 2000 and 2006, extent of habitat quality reduction, severity of habitat quality reduction and the product of area loss and the severity of habitat quality loss. Each point is a biotope, coloured according to the higher biotope classification. Point size is proportional to the number of species in each biotope.

When estimating diversity changes from the community specialization index (CSI) we find again that habitat quality changes better predict CSI changes than habitat quantity changes (Fig. 4). Across all habitats, increasing compound index area × severity habitat quality changes correlates with increasing loss of specialization both in 50 and 100 years. Increasing area with habitat quality loss also predicts CSI changes in 100 years (Supplementary Information S7).

**Figure 4.**
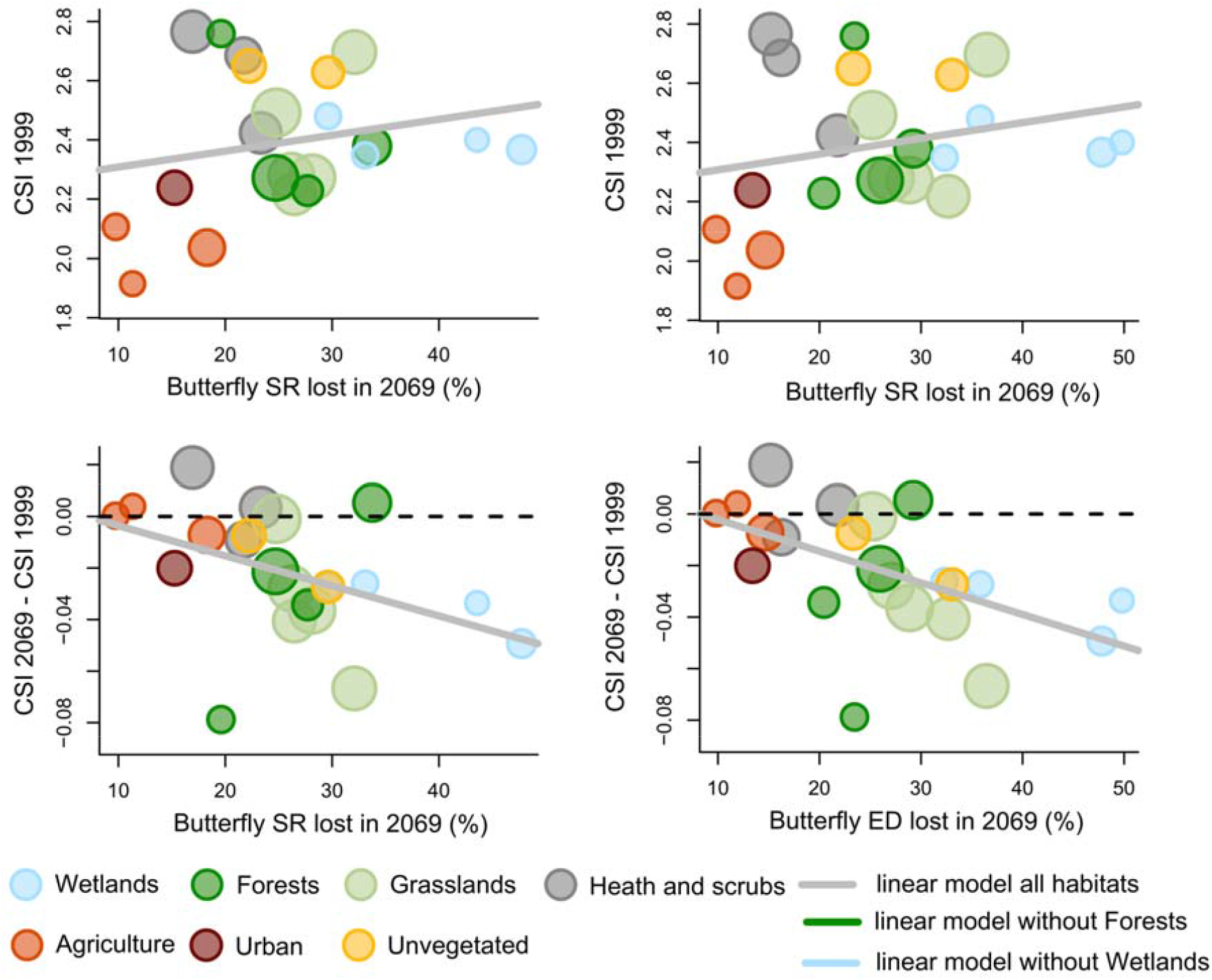
Relationship between CSI estimated before projections (1999), CSI change in 2069 (CSI 2069 minus CSI 1999) and predicted species richness (SR) and evolutionary distinctiveness (ED) lost in 2069. Each point is a biotope, coloured according to the higher biotope classification. Point size is proportional to the number of species in each biotope.

We do not find the community specialization index score to ever correlate with SR or ED loss in 2069 or 2119. However, we find a relationship between SR loss, ED loss and CSI changes at all periods of time (Supplementary Information S6-S7). Habitats with SR and ED declining faster are also habitats with faster decline of specialization.

Across all 23 habitats, both SR and ED changes in 2069 and 2119 are correlated with total area (Supplementary Information S6). The decline in SR and ED happens faster in smaller habitats and is accompanied by a faster loss of specialization of the habitats. Neither CSI scores nor CSI changes are significantly correlated with total area.

## Discussion

### The future of European butterflies

Population trends are difficult to estimate with the same precision for all species, even in Europe. The absolute quantification of biodiversity loss is therefore highly sensitive to the uncertainty in these estimates. However, the relative contribution of different habitats to diversity changes can still be estimated with more reliability from these population trend estimates, and we focus our discussion primarily on the comparison between habitats. Our projections also assumed that all conditions remained identical through time. How more complex models would affect our results is unclear. On the one hand, improvements of conservation practices or adaptive response from butterfly populations for example could clearly reduce the rate of decline. But on the other hand, other parameters such as climate change can both increase the rate of decline and extirpation of certain species while enabling others to thrive (Warren et al. 2021, Hill et al. 2021).

Concomitantly with species richness decline, we find that the relative proportion of habitat specialists is decreasing in comparison with generalists. We find that many habitats with the highest proportion of specialists are also among the most affected by this decline in specialization (e.g. Sclerophylous Shrubs, Broadleaved Evergreen Woodlands). Because specialists are projected to decline proportionally faster than generalists (Fig 4), future butterfly communities will increasingly consist of more homogeneous species assemblages, dominated by generalist species. Biotic homogenization driven by a disproportional loss of habitat specialists in butterflies results from many factors such as urbanization, habitat loss and climate change (Kuussaari et al. 2021, Warren et al. 2001, Ekroos et al. 2010). Homogenization is nowadays well characterized and considered a major trend in biodiversity responses to global changes, not only in butterflies but other groups as well such as ants (e.g. Hansen et al. 20 20), birds (e.g., Devictor et al. 2008, see also Bernardo-Madrid et al. 2019), or plants (e.g., Finderup Nielsen et al. 2019).

### Total habitat area but not cover changes predicts diversity loss

Total area currently covered by habitats is a strong predictor of SR and ED loss. Butterfly diversity in smaller habitats is declining faster and total area explains up to 41% of SR loss variance when excluding forest habitats. The effects of habitat fragmentation on diversity dynamics most likely increase in smaller habitats with smaller patch area, lower connectivity between patches and greater edge/interior ratio. These effects are known to negatively affect the metapopulations. For example, small patches prevent long-term persistence of local populations and increase the probability of emigration from patches (e.g. Sutcliffe et al.

1997). Haddad et al. (2015) experimentally showed that habitat fragmentation had both short-and long-term effects on the whole ecosystem, changing vegetation composition and successions, predation and herbivory. Fragmentation effects are also probably exacerbated in species with low dispersal capacity, a trait known to be correlated with faster rates of decline in butterflies (Kotiaho et al., 2005). Edge effects may also become increasingly negative when the edge/area ratio increases (Öckinger et al. 2012), in particular in landscapes with high density of arable lands where the use of fertilizers and pesticides affect the surrounding patches.

We find no relationship between changes in habitat cover over the last 50 years or between 2000 and 2006 and projected diversity changes. Habitat loss is considered as the major driver of insect decline, mediated by agricultural intensification and changes in farming practices (Warren et al. 2021) but our results do not directly support this observation. Given the spatial scale of our study, we cannot exclude that the effects of changes in habitat cover vary between habitats and across the geographical range considered here, hence distorting any relationship at the continental level. The fundamental components obscuring the relationship between habitat loss and diversity have been previously discussed for grassland butterflies (Sang et al. 2010). There is typically a lag between environmental changes and diversity response (extinction debt) and strong differences in sensitivity to habitat changes between habitat specialists and generalists. Both effects are likely to interact in our analyses and affect any relationship between habitat cover changes and diversity loss. In their long-term experiments, Haddad et al. (2015) reported long-lasting and delayed effects (up to 12 years) of habitat fragmentation on the decline in plants, birds and arthropods species richness.

### Total area of degraded habitats predicts diversity loss, except for forest habitats

Across all habitat types we find that the area of degraded habitats predicts the loss of specialist species. We also find strong evidence for a relationship between habitat quality changes and projected SR and ED loss, but this is not consistent across all habitat types. When excluding all forest habitats, the projected decline of butterflies accentuates in habitats with greater area affected by habitat quality loss. The four forest habitats included in our analysis all had low predicted biodiversity loss, considering that they scored the highest percentage of habitat characterized by quality loss. The reason behind this is unclear. The area of planted forest has only increased, at least since 1990 according to the FAO 2020, both worldwide and in Europe. Tree plantations in Europe in particular are often monocultures and composed of about 80% of exotic species (FAO 2020). Such plantations and intensively managed forests in general, most likely contribute to the high scores of quality degradation we find here. Despite that, butterfly decline is proportionally low. In our analyses, forests alongside with agricultural habitats were the most widespread habitats. This large network of forest habitats may help buffering to a certain extent the effects of quality degradation. However, strong differences may also exist geographically and between the different forest types, obscuring the relationship with diversity decline and more work is needed to understand population trends in forest habitat.

Our results indicate that wetland biotopes are characterized by both highest loss of diversity and widespread loss of habitat quality. Wetlands have long been recognized as highly endangered habitats and drastic declines of wetland fauna have been reported. For example, Fraixedas et al. (2017) estimated the rate of decline of peatland bird specialists in Finland to reach 2% every year since the 1980’s. Wetlands clearly stand out in our results with the strong relationship between the severity of habitat quality degradation and the decline of butterflies. We show that widespread decline in wetland habitat quality, rather than the intensity of that decline, best explains the great decline of butterflies. Wetlands have disappeared at a dramatically high rate over the 20^th^ century (Davidson 2014) and are currently threatened by drainage, direct exploitation and eutrophication. According to Reis et al. (2017), about 14.9% of wetlands are protected in Europe, of which only half are under strict protection. Our results suggest that increasing levels of protection and restoring for better quality wetlands, rather than maximizing land cover of low-quality habitats, is paramount for conservation of wetlands butterflies.

### Severity of habitat quality loss predicts diversity loss but for wetland habitats

Besides showing that area of habitat quality loss predicts biodiversity changes (Fig. 1,2), we also identify a strong correlation between the severity of habitat quality degradation and the decline in diversity, but only when wetlands are removed from our analyses. Grassland and Forest habitats in particular show both high intensity of habitat degradation, great species and phylogenetic diversity loss and decline of specialist species.

Grasslands are the most butterfly-species rich habitat in Europe and are known to be subject to a large number of threats. On the one hand, the changes in plant community composition and structure and associated modifications of the microclimate are detrimental to butterfly species that usually thrive in traditionally maintained, low-productivity agricultural landscapes (e.g. Habel et al. 2016). On the other hand, eutrophication, landscape homogenisation or changing land management that accompany agricultural intensification threaten oligotrophic habitat specialists, which either cannot find suitable host-plants or complete their life cycle (e.g. Wallisdevries et al. 2012, Pöyry et al. 2017).

### Prioritizing habitat quantity or habitat quality?

Overall, our results support the role of habitat quality changes (both area of quality loss and severity of quality loss) in predicting the decline of butterflies – both in diversity and specialization – much better than land cover changes. Above all other factors, the ability of butterflies to persist in a given habitat is bound by host-plant availability. The vegetation thriving in a degraded habitat is most likely uncharacteristic of the plant community of the same healthy habitat. Changes in vegetation composition and structure resulting from eutrophication for example are well-characterized (De Vries et al. 2007) and affect both host-plant communities and nectar availability. Host-plant availability, in turn, directly affects the probability of a butterfly population to maintain over generations. In that regard, host-plant specialist butterflies are probably the most sensitive to habitat degradation.

Evidence of this direct relationship can be found at the population level of individual species (e.g. Brunbjerg et al. 2017), communities (e.g. Stefanescu et al. (2009), and regional landscape (Habel et al. 2015). Curtis et al. (2015) for example showed that changes in host-plant abundance was a strong predictor of abundance for 27 species of butterflies across a regional landscape. The abundance and composition of plant communities providing nectar are also known to be strong determinants of population persistence in habitat patches (e.g. Fleishman et al. 2002, Wallisdevries et al. 2012). Taken together, our results show that butterfly biodiversity conservation cannot be envisioned without a simultaneous conservation of the vegetation communities, which provide suitable microclimatic conditions, host-plants and nectar resources.

Alongside the urge to expand the protection of land and oceans in the post-2020 Global Biodiversity Framework, the United Nations have declared 2021-2030 the Decade on Ecosystem Restoration. The focus of habitat restoration is largely targeted at restoring vegetation (Hale et al. 2019), but habitat restoration itself encapsulates a wide array of practices (Miller & Hobbs 2007). Our results indicate that neither habitat protection, nor habitat restoration, should focus solely on the maximization of habitat area. Not only countries have failed at reaching the current Convention on Biological Diversity target of 17% of terrestrial realms protected areas (Watson et al. 2014, UNEP-WCMC and IUCN Protected Plant: The World Database on Protected Areas https://www.protectedplanet.net/en) but evidence of “perverse” effects of such strategies have emerged (e.g. Barnes et al. 2018, Woodley et al. 2019, Visconti et al. 2019). The effectiveness of these protected areas in particular are being questioned, with evidence for example that protected areas are not located in important places for biodiversity (Venter et al. 2017) and not efficiently managed (Watson et al. 2014, Woodley et al. 2019). In the light of accumulating evidence, it is clear that habitat protection must move beyond area-based target and focus on restoring ecosystems to increase the effectiveness of protection measures and to provide the full range of ecological requirements for biodiversity (di Sacco et al. 2021).

## Acknowledgements

The authors declare no conflict of interest. We would like to thank Mariana García Criado for helping us obtain the data from the European Red List of Habitats report. The research presented in this paper is a contribution to the strategic research area Biodiversity and Ecosystems in a Changing Climate, BECC, which supported NC’s research work. We thank additional funding sources from the Swedish Research Council to SF, CBD and AA; and the Swedish Foundation for Strategic Research and the Royal Botanic Gardens, Kew to AA.

## References

Andermann, T., Faurby, S., Turvey, S. T., Antonelli, A., & Silvestro, D. (2020). The past and future human impact on mammalian diversity. Science advances, 6(36), eabb2313.

Barnes, M. D., Glew, L., Wyborn, C., & Craigie, I. D. (2018). Prevent perverse outcomes from global protected area policy. Nature Ecology & Evolution, 2(5), 759–762.

Bernardo-Madrid, R., Calatayud, J., González-Suárez, M., Rosvall, M., Lucas, P. M., Rueda, M., Antonelli, A., & Revilla, E. (2019). Human activity is altering the world’s zoogeographical regions. Ecology Letters, 22(8): 1297–1305.

Bossard, M., Feranec, J., Otahel, J., & Steenmans, C. (2000). The revised and supplemented Corine land cover nomenclature. European environment agency, Copenhagen.

Brunbjerg, A. K., Høye, T. T., Eskildsen, A., Nygaard, B., Damgaard, C. F., & Ejrnæs, R. (2017). The collapse of marsh fritillary (Euphydryas aurinia) populations associated with declining host plant abundance. Biological Conservation, 211, 117–124.

Ceballos, G., Ehrlich, P. R., Barnosky, A. D., García, A., Pringle, R. M., & Palmer, T. M. (2015). Accelerated modern human–induced species losses: Entering the sixth mass extinction. Science advances, 1(5), e1400253.

Curtis, R. J., Brereton, T. M., Dennis, R. L., Carbone, C., & Isaac, N. J. (2015). Butterfly abundance is determined by food availability and is mediated by species traits. Journal of Applied Ecology, 52(6), 1676–1684.

Davidson, N. (2014). How much wetland has the world lost? Long-term and recent trends in global wetland area. Marine and Freshwater Research, 65: 934–941.

Dennis, R. L., & Eales, H. T. (1997). Patch occupancy in Coenonympha tullia (Muller, 1764) (Lepidoptera: Satyrinae): habitat quality matters as much as patch size and isolation. Journal of Insect Conservation, 1(3), 167–176.

Devictor, V., Julliard, R., Clavel, J., Jiguet, F., Lee, A., & Couvet, D. (2008). Functional biotic homogenization of bird communities in disturbed landscapes. Global ecology and biogeography, 17(2), 252–261.

De Vries, W., Kros, H., Reinds, G. J., Wamelink, W., Mol, J., van Dobben, H., Bobbink, R., Emmett, B., Smart, S., Evans, C., Schlutow, A., Kraft, P., Belyazid, S., Sverdrup, H., van Hinsberg, A., Posch, M., & Hettelingh, J. P. (2007). Developments in deriving critical limits and modelling critical loads of nitrogen for terrestrial ecosystems in Europe. (No. 1382). Alterra.

Di Sacco, A., Hardwick, K., Blakesley, D., Brancalion, P. H., Breman, E., Rebola, L. C., Chomba, S., Dixon, K., Elliott, S., Ruyonga, G., Shaw, K., Smith, P., Smith, R. J., & Antonelli, A. (2021). Ten Golden Rules for Reforestation to Optimise Carbon Sequestration, Biodiversity Recovery and Livelihood Benefits. Global Change Biology, 27(7), 1328–1348.

Ekroos, J., Heliölä, J., & Kuussaari, M. (2010). Homogenization of lepidopteran communities in intensively cultivated agricultural landscapes. Journal of Applied Ecology, 47, 459–467.

Eskildsen, A., Carvalheiro, L. G., Kisslning, W. D., Biesmeijer, J. C., Schweiger, O., & Hoye, T. (2015). Ecological specialization matters: long-term trends in butterfly species richness and assemblage composition depend on multiple functional traits. Diversity and Distributions, 21, 792–802.

FAO and UNEP. 2020. The State of the World’s Forests 2020. Forests, biodiversity and people. Rome.

Filz, K. J., Engler, J. O., Stoffels, J., Weitzel, M., & Schmitt, T. (2013). Missing the target? A critical view on butterfly conservation efforts on calcareous grasslands in south-western Germany. Biodiversity and Conservation, 22(10), 2223–2241.

Finderup Nielsen, T., Sand-Jensen, K., Dornelas, M., & Bruun, H. H. (2019). More is less: net gain in species richness, but biotic homogenization over 140 years. Ecology Letters, 22(10), 1650–1657.

Fleishman, E., Ray, C., Sjögren-Gulve, P., Boggs, C. L., & Murphy, D. D. (2002). Assessing the roles of patch quality, area, and isolation in predicting metapopulation dynamics. Conservation Biology, 16(3), 706–716.

Fraixedas, S., Lindén, A., Meller, K., Lindström, Å., Keišs, O., Kålås, J. A., Husby, M., Leivits, A., Leivits, M., & Lehikoinen, A. (2017). Substantial decline of Northern European peatland bird populations: Consequences of drainage. Biological conservation, 214, 223–232.

Fridley, J. D., Vandermast, D. B., Kuppinger, D. M., Manthey, M., & Peet, R. K. (2007). Co-occurrence based assessment of habitat generalists and specialists: A new approach for the measurement of niche width. Journal of ecology, 95(4), 707–722.

Gilpin, M. (Ed.). (2012). Metapopulation dynamics: empirical and theoretical investigations. Academic press.

Habel, J. C., Segerer, A., Ulrich, W., Torchyk, O., Weisser, W. W., & Schmitt, T. (2016). Butterfly community shifts over two centuries. Conservation Biology, 30(4), 754–762.

Haddad, N. M., Brudvig, L. A., Clobert, J., Davies, K. F., Gonzalez, A., Holt, R. D., Lovejoy, T. E., Sexton J. O., Austin, M. P., Collins, C. D., Cook, W. M., Damschen E. I., Ewers, R. M., Foster, B. L., Jenkins, C. N., King, A J., Laurance, W. F., Levey, D. J., Margules, C. R., Melbourne, B. A., Nicholls, A. O., Orrock, J. L., Song D-X & Townshend, J. R. (2015). Habitat fragmentation and its lasting impact on Earth’s ecosystems. Science advances, 1(2), e1500052.

Hallmann, C. A., Sorg, M., Jongejans, E., Siepel, H., Hofland, N., Schwan, H., Stenmans, W., Müller, A., Sumser, H., Hörren, T., Goulson, D., & de Kroon, H. (2017). More than 75 percent decline over 27 years in total flying insect biomass in protected areas. PloS one, 12(10), e0185809.

Hansen, R. R., Nielsen, K. E., Offenberg, J., Damgaard, C., Byriel, D. B., Schmidt, I. K., Sørensen, P. B., Kjær, C. and Strandberg, M. T. (2020). Implications of heathland management for ant species composition and diversity–Is heathland management causing biotic homogenization? Biological Conservation, 242, p.108422.

Hanski, I., & Gilpin, M. (1991). Metapopulation dynamics: brief history and conceptual domain. Biological journal of the Linnean Society, 42(1-2), 3–16.

Hanski, I., & Thomas, C. D. (1994). Metapopulation dynamics and conservation: a spatially explicit model applied to butterflies. Biological Conservation, 68(2), 167–180.

Hale, R., Mac Nally, R., Blumstein, D. T., & Swearer, S. E. (2019). Evaluating where and how habitat restoration is undertaken for animals. Restoration Ecology, 27(4), 775–781.

Hill, G. M., Kawahara, A. Y., Daniels, J. C., Bateman, C. C., & Scheffers, B. R. (2021). Climate change effects on animal ecology: butterflies and moths as a case study. Biological Reviews, 96(5), 2113–2126.

IPBES (2019): Global assessment report on biodiversity and ecosystem services of the Intergovernmental Science-Policy Platform on Biodiversity and Ecosystem Services. E. S. Brondizio, J. Settele, S. Díaz, and H. T. Ngo (editors). IPBES secretariat, Bonn, Germany. 1148 pages.

Isaac, N. J., Turvey, S. T., Collen, B., Waterman, C., & Baillie, J. E. (2007). Mammals on the EDGE: conservation priorities based on threat and phylogeny. PloS one, 2(3), e296.

Janssen, J. A. M., Rodwell, J. S., García Criado, M., Arts, G. H. P., Bijlsma, R. J., & Schaminee, J. H. J. (2016) European red list of habitats: Part 2. Terrestrial and freshwater habitats. European Union.

Kotiaho, J. S., Kaitala, V., Komonen, A., & Päivinen, J. (2005). Predicting the risk of extinction from shared ecological characteristics. Proceedings of the National Academy of Sciences, 102(6), 1963–1967.

Kuussaari, M., Heliölä, J., Pöyry, J., & Saarinen, K. (2007). Contrasting trends of butterfly species preferring semi-natural grasslands, field margins and forest edges in northern Europe. Journal of Insect Conservation, 11(4), 351–366.

Kuussaari, M., Toivonen, M., Heliölä, J., Pöyry, J., Mellado, J., Ekroos, J., Hyyryläinen, V., Vähä-Piikkiö, I., Tiainen, J. (2021) Butterfly species’ responses to urbanization: differing effects of human population density and built-up area. Urban Ecosystems, 24(3), 515—27.

Macgregor, C. J., Williams, J. H., Bell, J. R., & Thomas, C. D. (2019). Moth biomass increases and decreases over 50 years in Britain. Nature Ecology & Evolution, 3(12), 1645– 1649.

Maes, D., & Van Dyck, H. (2001). Butterfly diversity loss in Flanders (north Belgium): Europe’s worst case scenario? Biological conservation, 99(3), 263–276.

Maxwell, S. L., Cazalis, V., Dudley, N., Hoffmann, M., Rodrigues, A. S., Stolton, S., Visconti, P., Woodley, S., Kingston, N., Lewis, E. & Maron, M. (2020). Area-based conservation in the twenty-first century. Nature, 586(7828), 217–227.

Miller, J. R., & Hobbs, R. J. (2007). Habitat restoration—Do we know what we’re doing? Restoration Ecology, 15(3), 382–390.

Moss, D., Wyatt B., Cornaert, M.H. & Roekaerts, M. (1991) CORINE Biotopes: the design, compilation and use of an inventory of sites of major importance for nature conservation in the European Community Office for Official Publications of the European Communities Luxembourg.

Neubauer, T. A., Hauffe, T., Silvestro, D., Schauer, J., Kadolsky, D., Wesselingh, F. P., … & Wilke, T. (2021). Current extinction rate in European freshwater gastropods greatly exceeds that of the late Cretaceous mass extinction. Communications Earth & Environment, 2(1), 1–7.

Obura, D. O., Katerere, Y., Mayet, M., Kaelo, D., Msweli, S., Mather, K., Harris, J., Louis, M., Kramer, R., Teferi, T. & Samoilys, M. (2021). Integrate biodiversity targets from local to global levels. Science, 373(6556), 746–748.

Öckinger, E., Bergman, K. O., Franzén, M., Kadlec, T., Krauss, J., Kuussaari, M., Juha Pöyry, Smith, H.G., Steffan-Dewenter, I., & Bommarco, R. (2012). The landscape matrix modifies the effect of habitat fragmentation in grassland butterflies. Landscape Ecology, 27(1), 121– 131.

Pöyry, J., Carvalheiro, L. G., Heikkinen, R. K., Kühn, I., Kuussaari, M., Schweiger, O., Valtonen, A., van Bodegom, P. M., & Franzén, M. (2017). The effects of soil eutrophication propagate to higher trophic levels. Global Ecology and Biogeography, 26(1), 18–30.

Reis, V., Hermoso, V., Hamilton, S. K., Ward, D., Fluet-Chouinard, E., Lehner, B., & Linke, S. (2017). A global assessment of inland wetland conservation status. Bioscience, 67(6), 523– 533.

Sang, A., Teder, T., Helm, A., & Pärtel, M. (2010). Indirect evidence for an extinction debt of grassland butterflies half century after habitat loss. Biological Conservation, 143(6), 1405– 1413.

Schlicht, D. W., & Orwig, T. T. (1998). The status of Iowa’s Lepidoptera. J Iowa Acad Sci 105, 82–88.

Seibold, S., Gossner, M. M., Simons, N. K., Blüthgen, N., Müller, J., Ambarl, D., Ammer, C., Bauhus, J., Fischer, M., Habel, J. C., Linsenmair, K. E., Nauss, T., Penone, C., Prati, D., Schall, P., Schulze, E.-D., Vogt, J., Wöllauer, S., & Weisser, W. W. (2019). Arthropod decline in grasslands and forests is associated with landscape-level drivers. Nature, 574(7780), 671–674.

Stefanescu, C., Penuelas, J., & Filella, I. (2009). Rapid changes in butterfly communities following the abandonment of grasslands: a case study. Insect Conservation and Diversity, 2(4), 261–269.

Sutcliffe, O. L., Thomas, C. D., & Peggie, D. ournal(1997). Area-dependent migration by ringlet butterflies generates a mixture of patchy population and metapopulation attributes. Oecologia, 109(2), 229–234.

van Swaay, C., Cuttelod, A., Collins, S., Maes, D., López Munguira, M., Šašic, M., Settele, J., Verovnik, R., Verstrael, T., Warren, M., Wiemers, M., & Wynhoff, I. (2010): European Red List of butterflies. (IUCN Red List of Threatened Species - Regional Assessment) - Office for Official Publications of the European Communities, Luxembourg

Thomas, J. A., Bourn, N. A. D., Clarke, R. T., Stewart, K. E., Simcox, D. J., Pearman, G. S., Curtis, R., & Goodger, B. (2001). The quality and isolation of habitat patches both determine where butterflies persist in fragmented landscapes. Proceedings of the Royal Society of London. Series B: Biological Sciences, 268(1478), 1791–1796.

van Strien, A. J., van Swaay, C. A., van Strien-van Liempt, W. T., Poot, M. J., & WallisDeVries, M. F. (2019). Over a century of data reveal more than 80% decline in butterflies in the Netherlands. Biological Conservation, 234, 116–122.

Van Swaay, C., Warren, M., & Loïs, G. (2006). Biotope use and trends of European butterflies. Journal of Insect Conservation, 10(2), 189–209.

Van Swaay Cam, Dennis EB, Sch mucki R, Sevilleja C, Balalaikins M, Botham M, Bourn N, Brereton T, Cancela JP, Carlisle B, Chambers P, Collins S, Dopagne C, Escobés R, Feldmann R, Fernández-García JM, Fontaine B, Gracianteparaluceta A, Harrower C, Harpke A, Heliölä J, Komac B, Kühn E, Lang A, Maes D, Mestdagh X, Middlebrook I, Monasterio Y, Munguira ML, Murray TE, Musche M, Õunap E, Paramo F, Pettersson LB, Piqueray J, Settele J, Stefanescu C, Švitra G, Tiitsaar A, Verovnik R, Warren MS, Wynhoff I, Roy DB (2019). The EU Butterfly Indicator for Grassland species: 1990-2017: Technical Report. Butterfly Conservation Europe.

Venter, O., Magrach, A., Outram, N., Klein, C. J., Possingham, H. P., Di Marco, M., & Watson, J. E. (2018). Bias in protected-area location and its effects on long-term aspirations of biodiversity conventions. Conservation Biology, 32(1), 127–134.

Visconti, P., Butchart, S. H., Brooks, T. M., Langhammer, P. F., Marnewick, D., Vergara, S., Yanosky, A. and Watson, J. E. (2019). Protected area targets post-2020. Science, 364(6437), 239–241.

Wagner, D. L. (2020). Insect declines in the Anthropocene. Annual review of entomology, 65, 457–480.

Wallisdevries, M. F., van Swaay, C. A., & Plate, C. L. (2012). Changes in nectar supply: a possible cause of widespread butterfly decline. Current Zoology, 58(3), 384–391.

Warren, M. S., Hill, J. K., Thomas, J. A., Asher, J., Fox, R., Huntley, B., Roy, D. B., Telfer, M. G., Jeffcoate, S., Harding, P., Jeffcoate, G., Willis, S. G., Greatorex-Davies, J. N., Moss, D., & Thomas, C. D. (2001). Rapid responses of British butterflies to opposing forces of climate and habitat change. Nature, 414(6859), 65–69.

Warren, M. S., Maes, D., van Swaay, C. A., Goffart, P., Van Dyck, H., Bourn, N. A., Wynhoff, I., Hoare, D., & Ellis, S. (2021). The decline of butterflies in Europe: Problems, significance, and possible solutions. Proceedings of the National Academy of Sciences, 118(2), e2002551117.

Watson, J. E., Dudley, N., Segan, D. B., & Hockings, M. (2014). The performance and potential of protected areas. Nature, 515(7525), 67–73.

Wiemers, M., Chazot, N., Wheat, C. W., Schweiger, O., & Wahlberg, N. (2020). A complete time-calibrated multi-gene phylogeny of the European butterflies. ZooKeys, 938, 97.

Wiemers, M., Balletto, E., Dincă, V., Fric, Z. F., Lamas, G., Lukhtanov, V., Munguira, M. L., van Swaay, C. A. M., Vila, R., Vliegenthart, A., Wahlberg, N. & Verovnik, R. 2018: An updated checklist of the European butterflies (Lepidoptera: Papilionoidea). ZooKeys 811: 9–45. doi:10.3897/zookeys.811.28712

Woodley, S., Locke, H., Laffoley, D., MacKinnon, K., Sandwith, T., & Smart, J. (2019). A review of evidence for area-based conservation targets for the post-2020 global biodiversity framework. Parks, 25(2), 31–46.

WWF (2020) Living Planet Report 2020.

