## Supplementary figures and images for "Large-scale assessment of habitat quality and quantity change on declining European butterflies"

### Supplementary Information S3.1

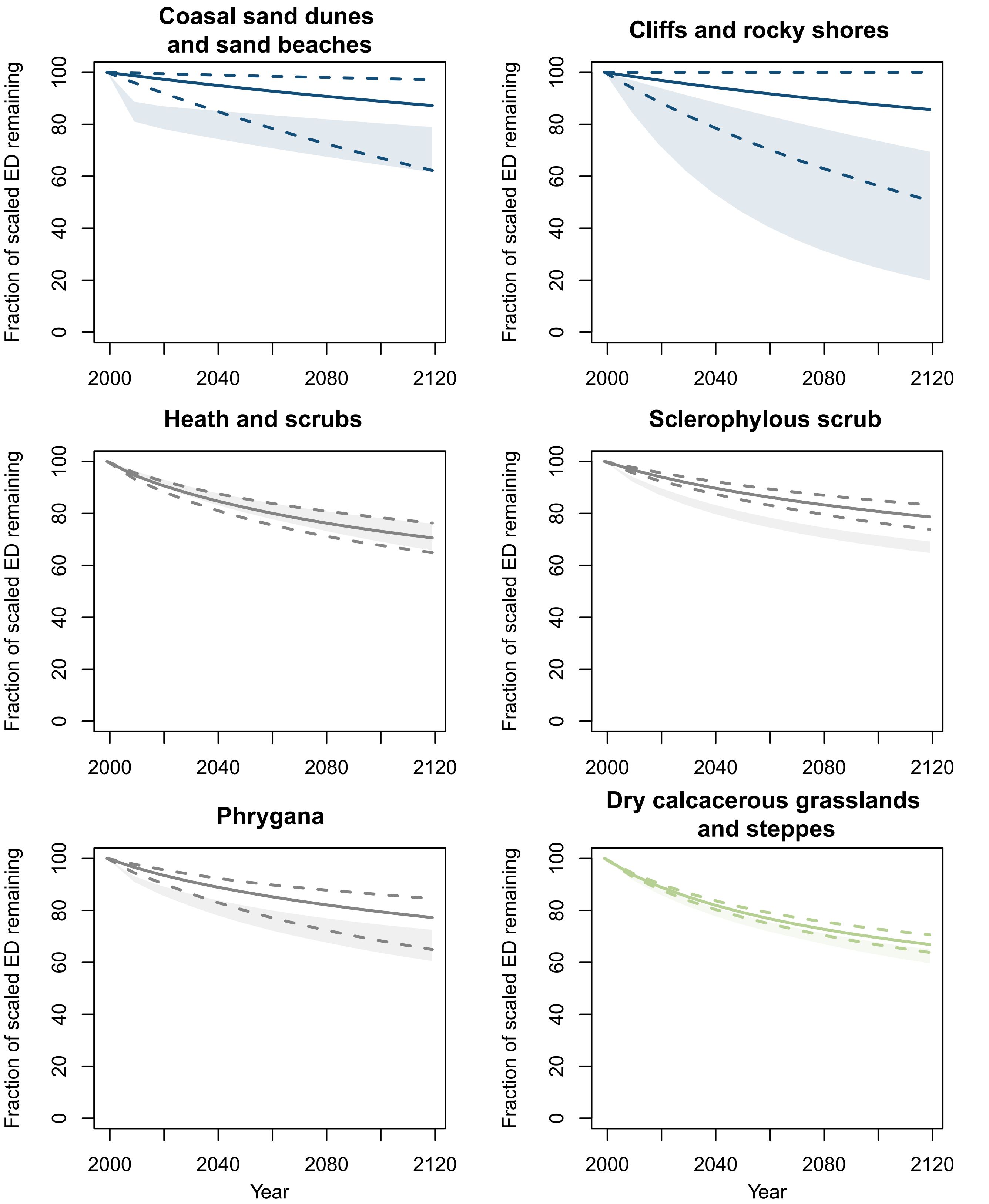

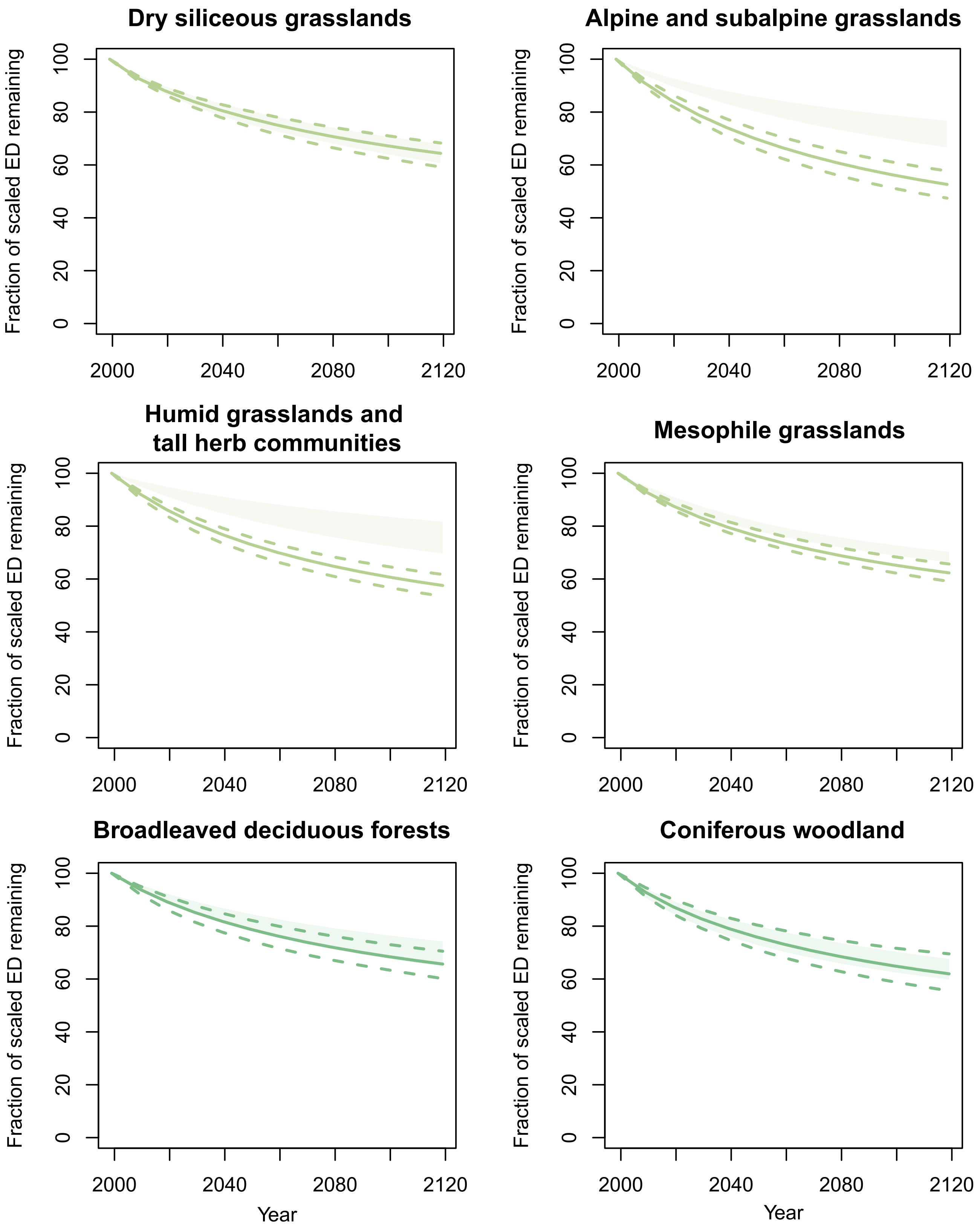

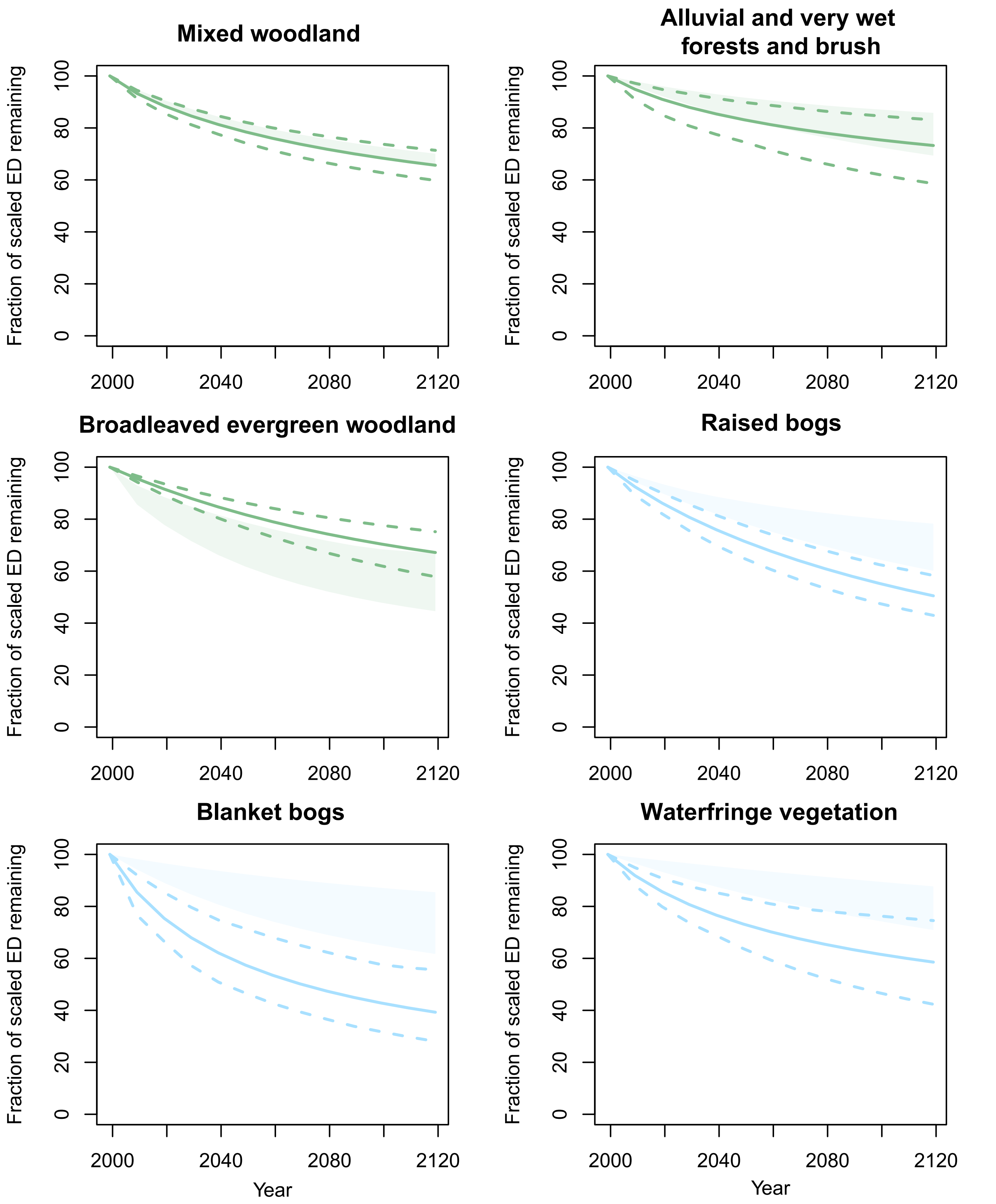

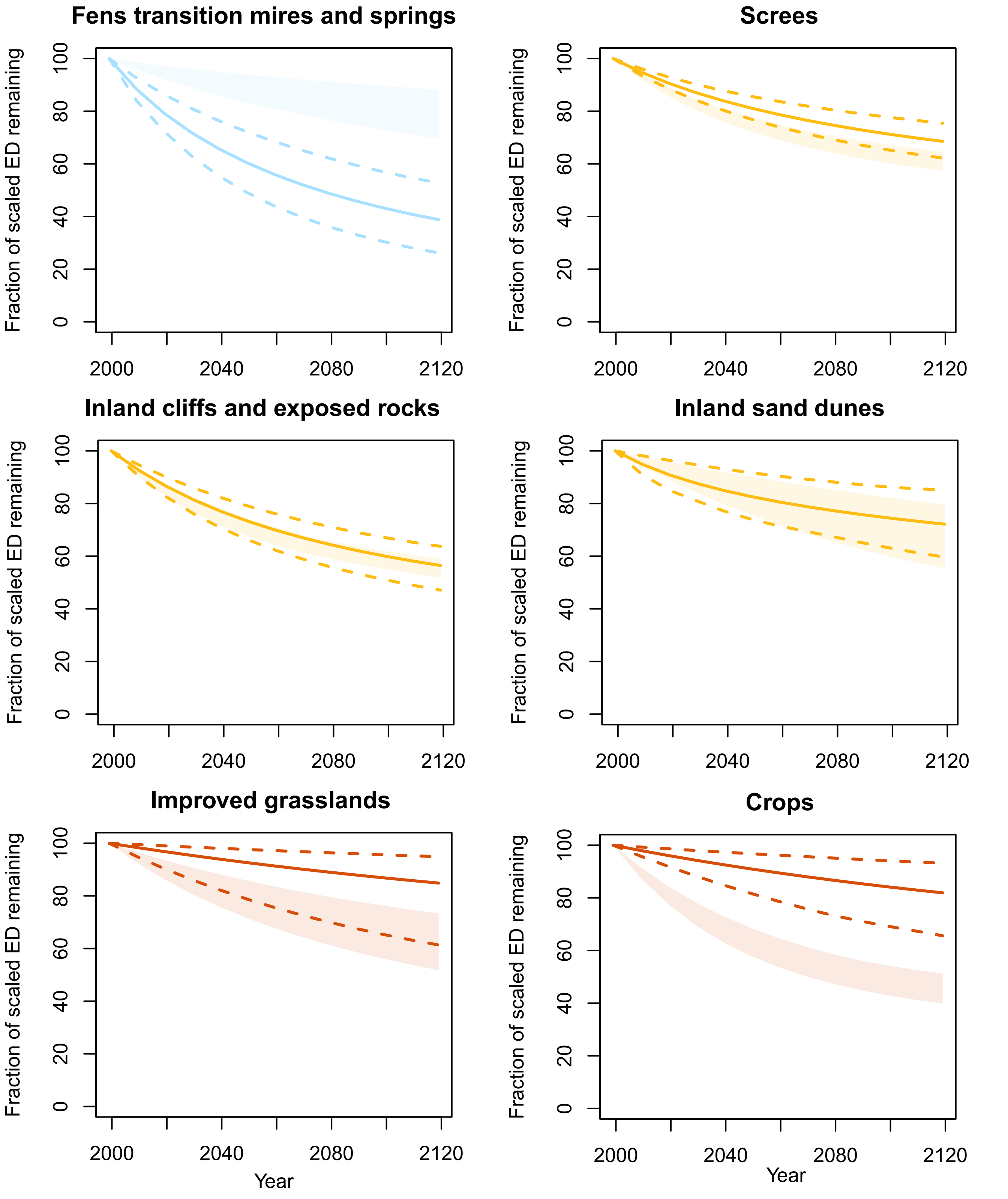

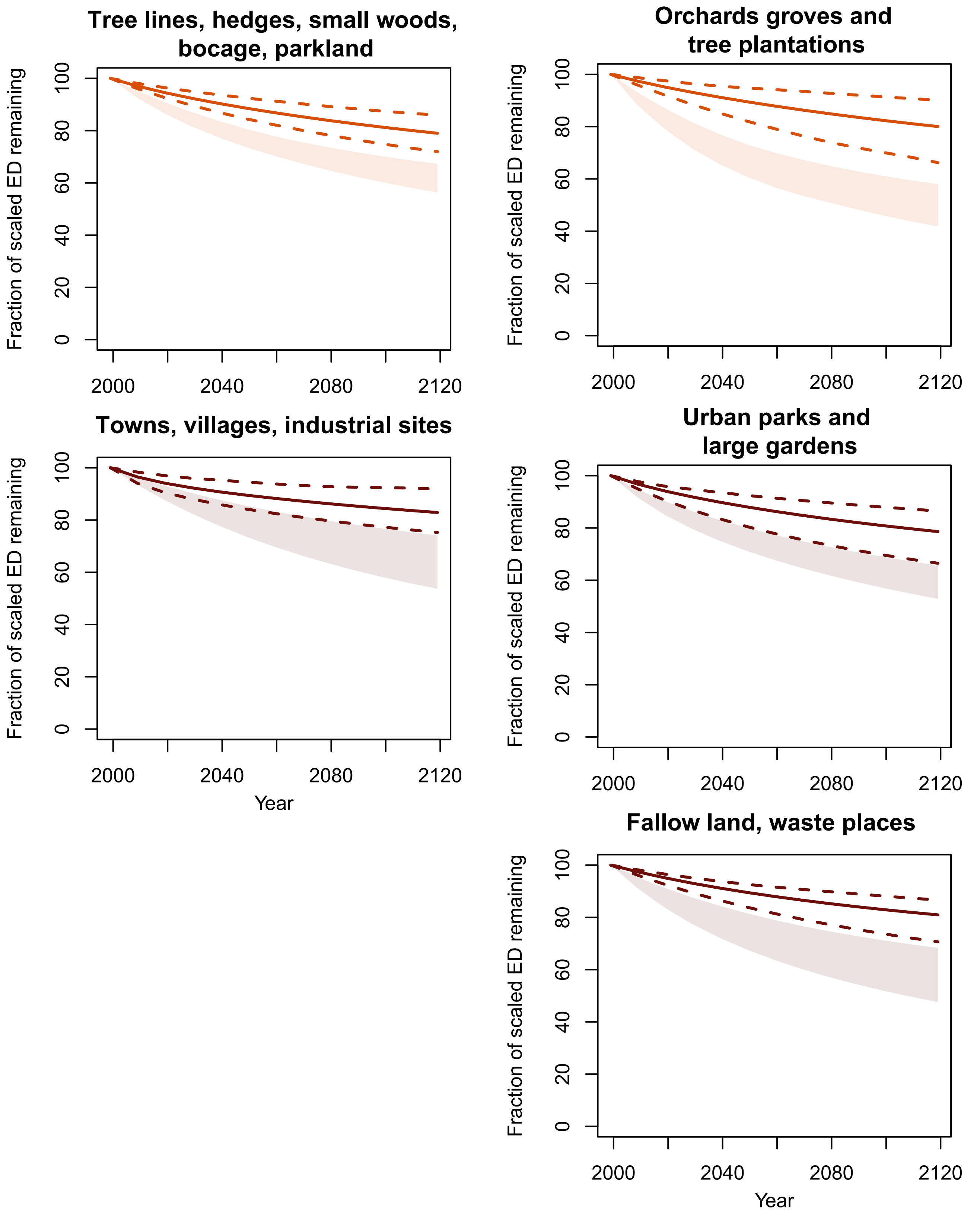

### Supplementary Information S3.2

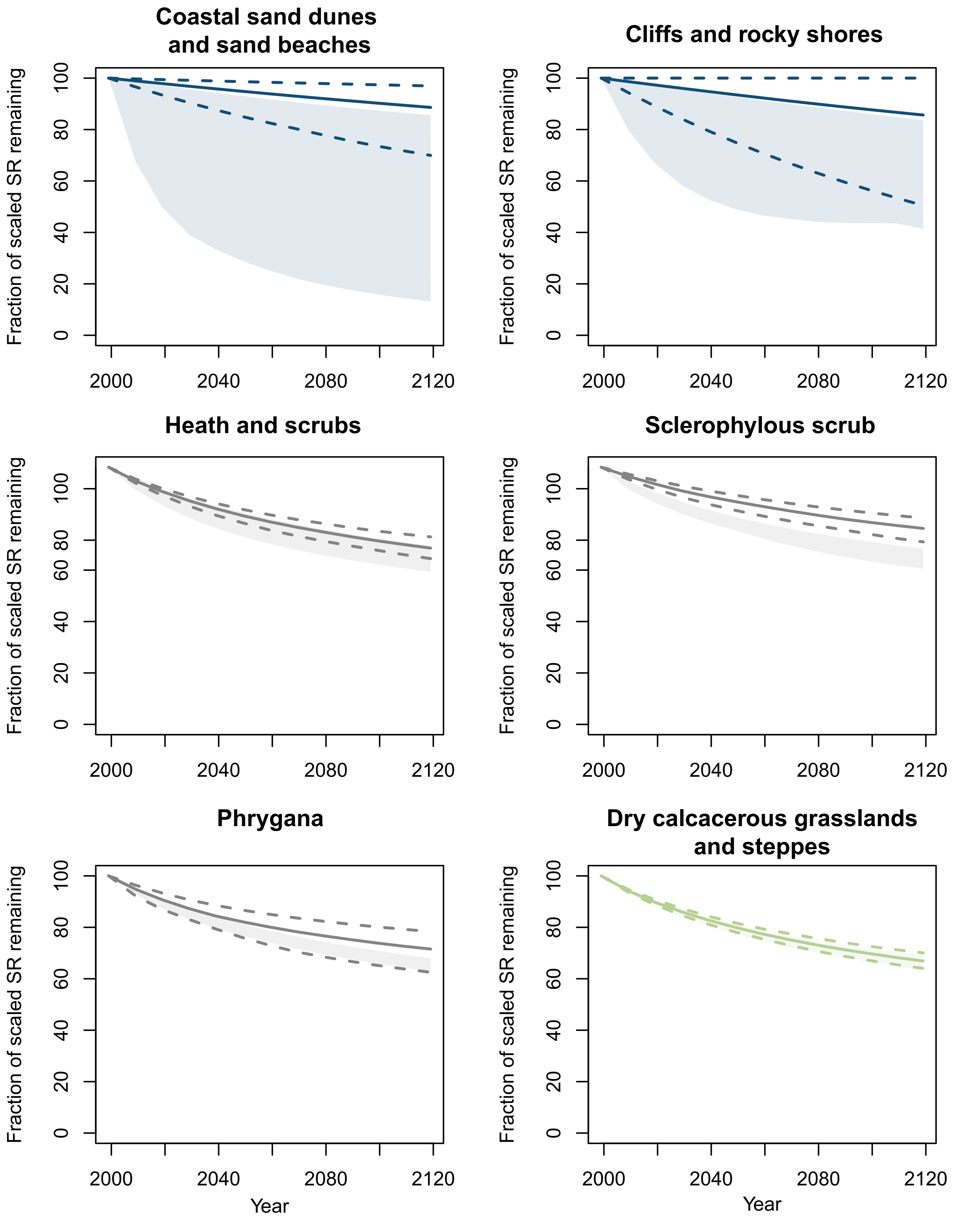

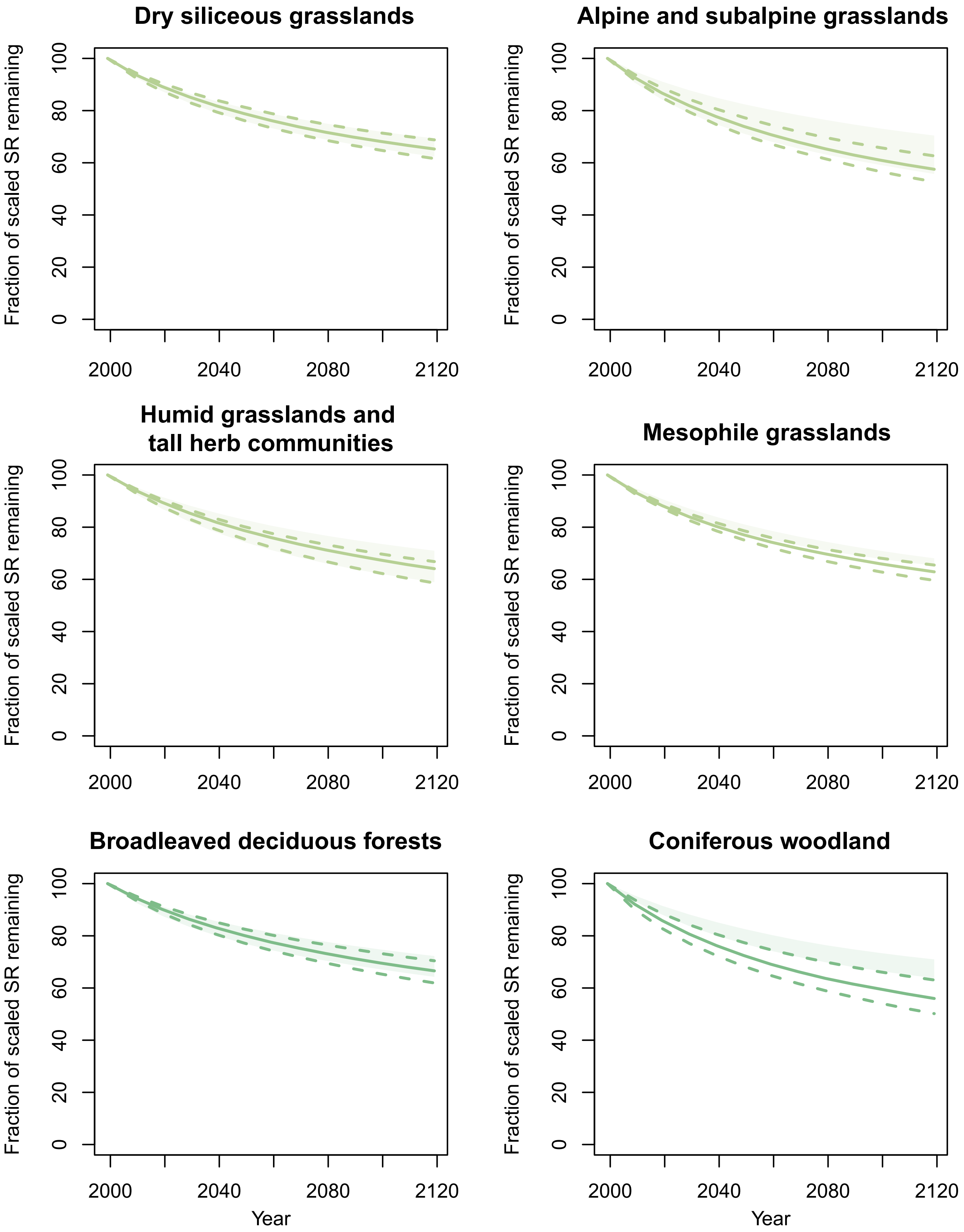

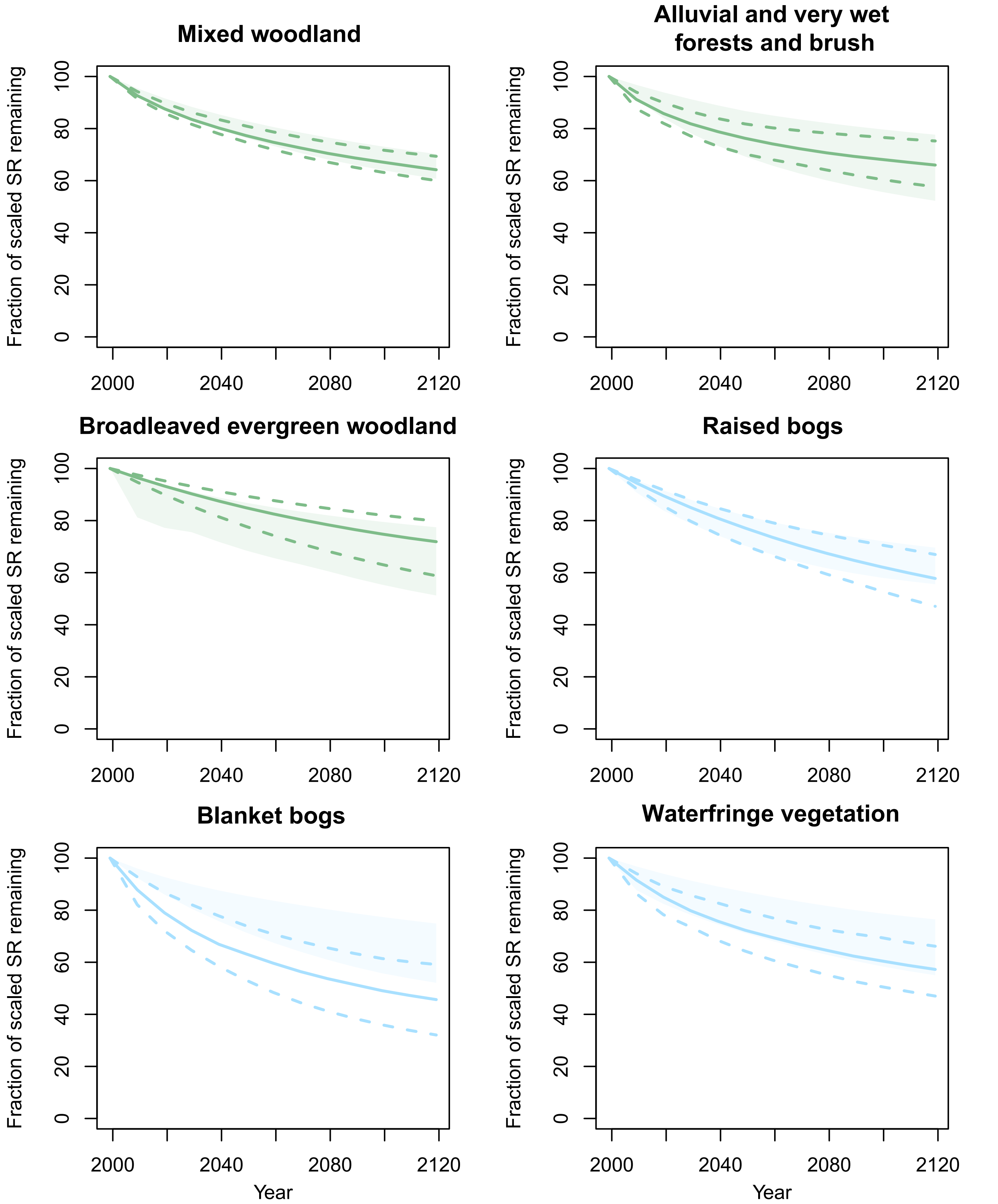

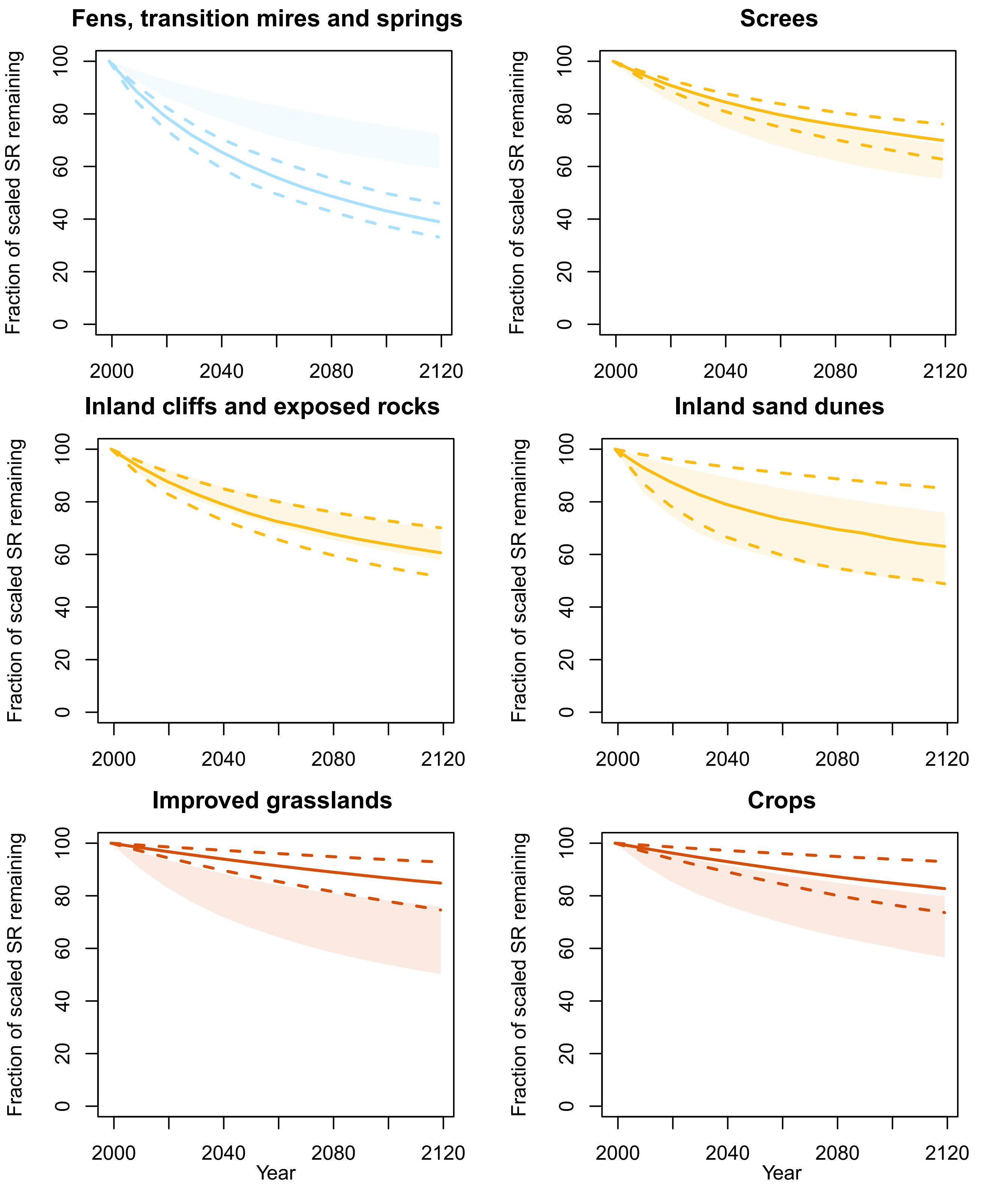

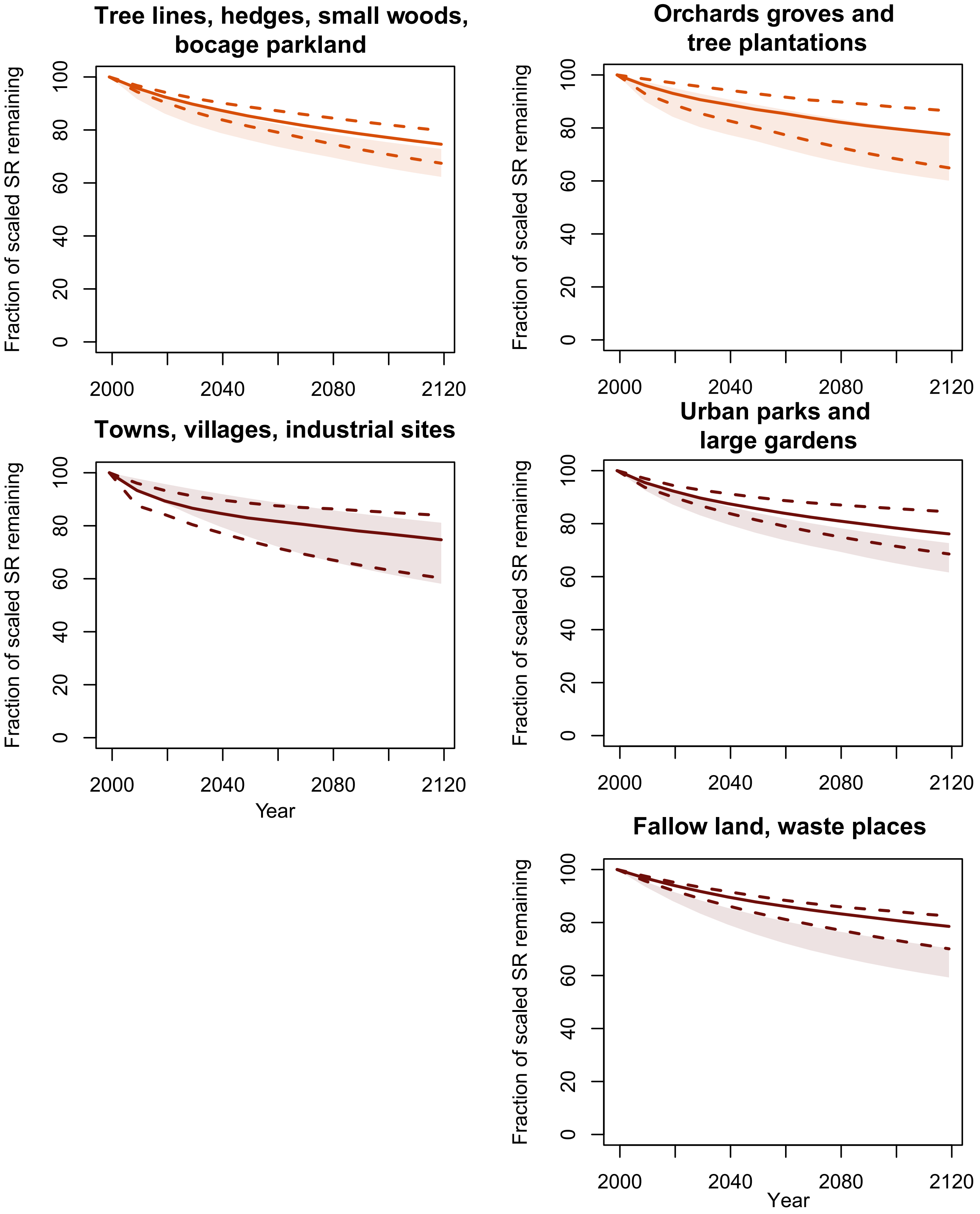
