## Supplementary Information for "Large-scale assessment of habitat quality and quantity change on declining European butterflies"

### 1- Dataset

Table S1.1. Population trends between 1999 and 2009 used to produce the 2009 report on the state of European butterflies (https://ec.europa.eu/environment/nature/conservation/species/redlist/downloads/European_butterflies.pdf).

Table S1.2. Habitat occupancy dataset used in this study. This dataset is modified from "van Swaay, C., Warren, M., & Loïs, G. (2006). Biotope use and trends of European butterflies. Journal of Insect Conservation, 10(2), 189-209." to match the species list from "Wiemers, M., Chazot, N., Wheat, C. W., Schweiger, O., & Wahlberg, N. (2020). A complete time-calibrated multi-gene phylogeny of the European butterflies. ZooKeys, 938, 97."

Table S1.3. Equivalence table between the classification used by "van Swaay, C., Warren, M., & Loïs, G. (2006). Biotope use and trends of European butterflies. *Journal of Insect Conservation*, 10(2), 189-209." and the Corine classification (CLC) as used by the Corine Land Cover European program (https://www.eea.europa.eu/data-and-maps/data/corine-biotopes).

Table S1.4. Equivalence table between EUNIS habitat classification and Corine classification (CLC) as used by the Corine Land Cover European program (https://www.eea.europa.eu/data-and-maps/data/corine-biotopes).

### 2- Weighted Species richness and Evolutionary Distinctiveness

At each step in the projection, species richness of each habitat was calculated as the sum of the remaining population sizes, weighted by habitat use. For example, in Figure S2 species richness of habitat 1 is calculated as (100+0.3*70+0.5*50). Hence, the total remaining population of each species is partitioned across habitat according to its relative habitat use.

To estimate phylogenetic diversity changes through time in different habitats we used the Evolutionary Distinctiveness index (ED). However, we also partitioned ED into different habitats proportionally to its relative use of these habitats. Thus, for a single species, the sum of ED scores across the habitats used by that species is equal to its standard ED score. As a result, a species contributed to the habitat total ED in proportion of its habitat use. In addition, a species contribution to the habitat total ED at a time t in our projections, was weighted by the proportion of the initial population remaining at that time.

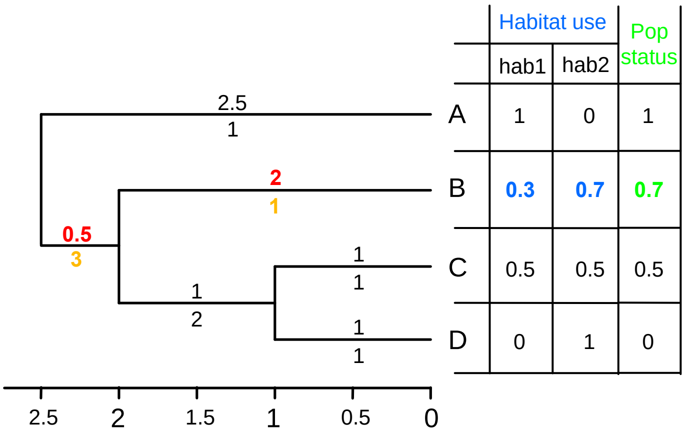


| | Habitat use | | Pop status |
|---|---|---|---|
| | hab1 | hab2 | |
| A | 1 | 0 | 1 |
| B | **0.3** | **0.7** | **0.7** |
| C | 0.5 | 0.5 | 0.5 |
| D | 0 | 1 | 0 |

$ED_{B\text{-}hab1} = (0.5/3)*0.3*0.7 + (2/1)*0.3*0.7$

$ED_{B\text{-}hab2} = (0.5/3)*0.7*0.7 + (2/1)*0.7*0.7$

Figure S2. Theoretical example of the calculation of the weighted ED. In this example we show how much ED species B contributes to habitat 1 (hab1) and habitat 2 (hab2) given its relative habitat use (blue numbers) and its population status at time t. For habitat 1, branch lengths are divided by the number of species descending from that branch (standard ED calculation) and multiplied by the relative frequency of habitat use and population status. The sum of these weighted branch lengths is the contribution of species B to habitat 1 total ED at a time t.

3- Results of projected SR and ED loss for every habitat

Figure S3.1 (separate document). Projected SR loss in the future for all habitats. Plain and dashed lines represent the median and 95% confidence interval respectively of the 1000 projections. Shaded area represents the 95% confidence interval of projected SR loss after 1000 permutations of population trends.

Figure S3.2 (separate document). Projected ED loss in the future for all habitats. Plain and dashed lines represent the median and 95% confidence interval respectively of the 100 projections, each permutation including a different phylogenetic tree sampled from Wiemers et al. (2020) posterior distribution. Shaded area represents the 95% confidence interval of projected ED loss after 100 permutations of population trends.

4- Changes in Community Specialization Index (CSI) in 50 and 100 years.

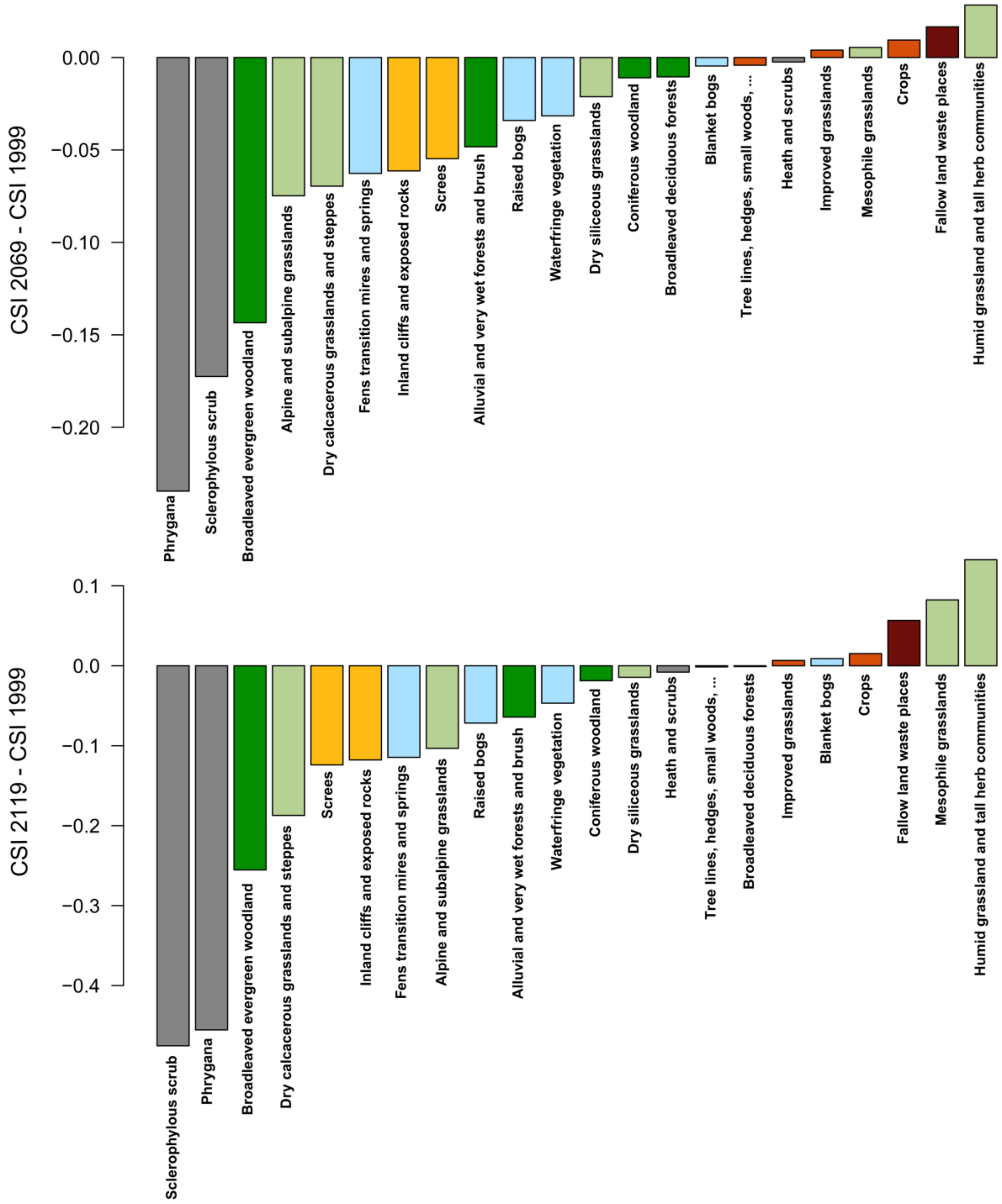


Figure S4. Changes in Community Specialization Index (CSI) in 50 years (top) and 100 years (bottom) for all habitats included in our analyses. Negative values indicate a loss of specialization, hence a proportionally higher decline of specialist species.

<u>5- Relationship between the predicted evolutionary distinctiveness (ED) lost in 2069 and habitat changes.</u>

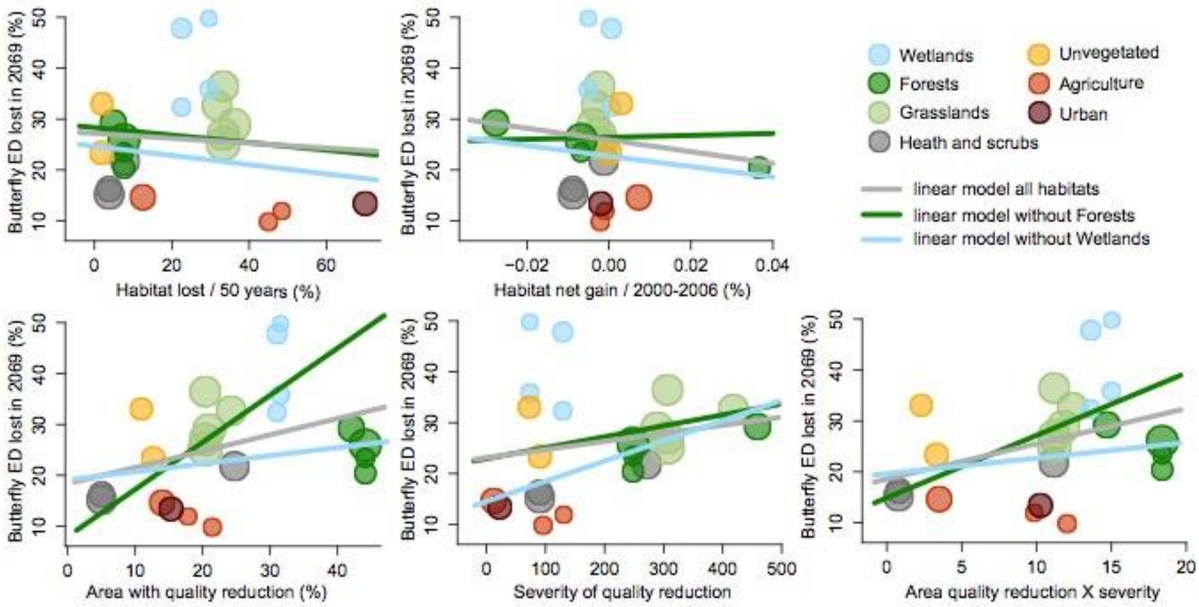


Figure S5. Relationship between the predicted evolutionary distinctiveness (ED) lost in 2069 and habitat net gain over the last 50 years, habitat net gain between 2000 and 2006, extent of habitat quality reduction, severity of habitat quality reduction and the product of area loss and the severity of habitat quality loss. Dot size is proportional to the number of species in each biotope.

6- Relationship between predicted diversity lost in 2069 and the log total habitat surface from the CLC dataset.

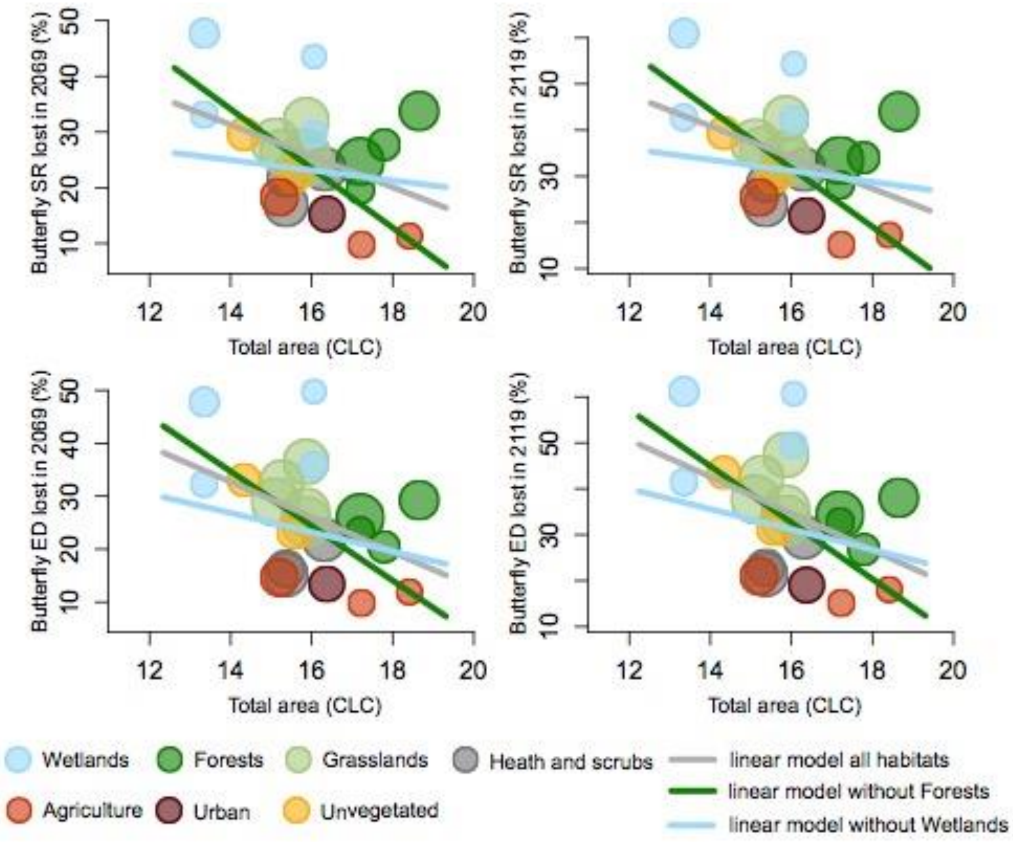

Figure S6. Relationship between predicted species richness (SR) and evolutionary distinctiveness (ED) lost in 2069 and the log total habitat surface from the CLC dataset

7- Results of linear models

Table S7.1. Results of linear models fitted between projected SR loss and the different estimation of habitat changes, and the habitat total area. Results include the models run after removing all Forest and Wetlands habitats. The likelihood (logLik), the AIC score (AIC), the slope, $R^2$ and pvalue of the model are reported.

Table S7.2. Results of linear models fitted between projected ED loss and the different estimation of habitat changes, and the habitat total area. Results include the models run after removing all Forest and Wetlands habitats. The likelihood (logLik), the AIC score (AIC), the slope, $R^2$ and pvalue of the model are reported.

Table S7.3. Results of linear models fitted between CSI score in 1999 and projected CSI score in 2069 and 2119, and the different estimation of habitat changes, and the habitat total area. Results include the models run after removing all Forest and Wetlands habitats. The likelihood (logLik), the AIC score (AIC), the slope, $R^2$ and pvalue of the model are reported.

Table S7.4. Results of linear models fitted between CSI difference between 2069 and 1999 and between 2119 and 1999, and the different estimation of habitat changes, and the habitat total area. Results include the models run after removing all Forest and Wetlands habitats. The likelihood (logLik), the AIC score (AIC), the slope, $R^2$ and pvalue of the model are reported.

Table S7.5. Results of linear models fitted between projected SR loss, ED loss, total habitat area, and CSI scores in 1999, 2069, 2119 and CSI difference between 2069 and 1999 and between 2119 and 1999.the habitat total area. Results include the models run after removing all Forest and Wetlands habitats. The likelihood (logLik), the AIC score (AIC), the slope, $R^2$ and pvalue of the model are reported.
